# A toolbox of Stable Integration Vectors (SIV) in the fission yeast *Schizosaccharomyces pombe*

**DOI:** 10.1101/808329

**Authors:** Aleksandar Vještica, Magdalena Marek, Pedro N’kosi, Laura Merlini, Gaowen Liu, Melvin Bérard, Ingrid Billault-Chaumartin, Sophie G Martin

## Abstract

*Schizosaccharomyces pombe* is a widely used model organism that resembles higher eukaryotes in many aspects of cell physiology. Its popularity as an experimental system partially stems from the ease of genetic manipulations, where the innate homology-targeted repair is exploited to precisely edit the genome. While vectors to incorporate exogenous sequences into the chromosomes are available, most are poorly characterized. Here we show that commonly used fission yeast vectors, which upon integration produce repetitive genomic regions, yield unstable genomic loci. We overcome this problem by designing a new series of Stable Integration Vectors (SIV) that target four different prototrophy genes. SIV produce non-repetitive, stable genomic loci and integrate predominantly as single copy. Additionally, we develop a set of complementary auxotrophic alleles that preclude false-positive integration events. We expand the vector series to include antibiotic resistance markers, promoters, fluorescent tags and terminators, and build a highly modular toolbox to introduce heterologous sequences. Finally, as proof of concept, we generate a large set of ready-to-use, fluorescent probes to mark organelles and cellular processes with a wide range of applications in fission yeast research.

## Introduction

The fission yeast *Schizosaccharomyces pombe* is a well-established model organism for studying diverse aspects of cellular biology. It continues to play a critical role for instance in the discovery of fundamental aspects of cell cycle control, chromosome biology, signaling and cytoskeleton dynamics (Hoffman et al., 2015). Decades of research have yielded versatile molecular methods to control the expression of native and foreign genes in this species.

The ability to transform cells with genetic material and create transgenic lines is critical in biological research. Foreign DNA fragments are easily delivered to *S. pombe* cells, where they have one of two fates. First, circular DNA plasmids can be maintained as episomal fragments, provided they contain an autonomous replicating sequence (ARS) (*e.g.* the pREP series; Craven et al., 1998; Forsburg and Sherman, 1997; Maundrell, 1993; Moreno et al., 2000). However, these episomal plasmids do not contain centromeric segments, because *S. pombe* centromeres are complex 100kb-long sequences, distinct from the point centromeres of the budding yeast *S. cerevisiae* (Clarke, 1990; Yamagishi et al., 2014). For this reason, such circular plasmids segregate randomly during division, leading to variable copy numbers in the population and the need for continuous selective pressure to prevent plasmid loss. Second, because homologous recombination is highly efficient, linear DNA fragments can easily integrate at desired genomic loci. This allows precise genome editing and direct gene manipulation at their native locus, the method of choice to alter gene function in near-physiological conditions. It also allows for integration of linearized ‘integrative’ plasmids, for instance containing foreign DNA, to placeholder genomic loci. The most common integrative vectors typically carry a single homology region. Plasmid linearization within this region enables the homology directed repair to target the vector to the desired genomic location (Keeney and Boeke, 1994; Matsuyama et al., 2004; Maundrell, 1993). However, this system suffers from the problem that integration leads to duplication of sequences on either side of the integrated vector (Siam et al., 2004), which can further recombine and lead to either amplification or deletion of integrated fragment. This is especially noticeable if the insert alters cell fitness, and can cause reproducibility issues between experiments. A few integrative plasmids that should not lead to the formation of genomic copies have been developed, but their stability has not been directly probed (Fennessy et al., 2014; Kakui et al., 2015).

Here, we present a series of easy-to-use, modular integrative vectors that insert into the genome as a stable, single copy, without formation of genomic repeats. The basic elements included in our vector series are: a bacterial replication origin and AmpR allowing for plasmid amplification in bacteria, one or two multicloning sites (MCS), a sequence targeting the construct to one of four different chromosomal locations (*ade6*, *ura4*, *lys3*, *his5*), which confers prototrophy post transformation, and an optional drug resistance marker. We further expand this basic vector backbone by including various promoter, fluorescence tag and terminator sequences, allowing for expression of any gene of interest at desired levels.

The most popular inducible promoter in fission yeast is based on the strong *nmt1* (**n**o **m**essage in **t**hiamine) promoter and its two attenuated versions that carry mutations in the TATA box – namely *nmt41* and *nmt81* (Basi et al., 1993; Maundrell, 1990; Maundrell, 1993). The *nmt* promoters display a strong induction fold after de-repression (estimated at about 80-fold for *nmt1*; Basi et al., 1993), but their maximal induction time exceeds 15h and is not completely synchronous in the population (Maundrell, 1990; Watson et al., 2013). The *urg1* promoter is also inducible and shows a strong induction in the presence of uracil (but also upon nitrogen starvation), with faster kinetics: maximal transcript levels occur 30 min post induction and decrease rapidly after uracil removal with both states being stable for at least 24h (Watt et al., 2008). However, at non-native integration sites, the off-state of *p^urg1^* is strongly elevated, reducing the dynamic range of induction (Watson et al., 2013). The most widely used constitutive promoter is the strong *p^adh1^*, and a few others of similar or slightly weaker strength have been described (Matsuyama et al., 2008; Siam et al., 2004). Here, we make use of the known inducible promoters (*nmt1*, *nmt41*, *nmt81*, *urg1*) and a series of constitutive promoters (*cdc12*, *pom1*, *rga3*, *pak1*, *act1* and *tdh1*) that lead to GFP expression over three orders of magnitude.

Our Stable Integration Vector (SIV) series was built in a highly modular way. For instance, fluorescence tagging vectors allow for both N-and C-terminal tagging of the construct of interest. As multicloning sites are identical in all the vectors, quick exchange of elements allows for further expansion of the toolbox. We demonstrate the stable and single-copy integration of the SIV series and fully describe promoter strength. We introduce three new auxotrophic deletion alleles of the target genomic integration sites (at *ade6*, *lys3*, *his5*), which abrogate false-positive transformants. Finally, using the SIV series, we generate a panel of fluorescent bio-markers in three compatible wavelengths (mTagBFP2, sfGFP or GFP and mCherry) labelling commonly studied cellular organelles, structures and processes. We expect these tools to be a valuable asset to all researchers using the fission yeast system.

## Results

### The presence of two regions of homology to target sequences promotes stability of integrants and avoids tandem integration events

In fission yeast, homologous recombination is efficient and has been used for decades to introduce exogenous constructs at defined genomic loci through integration of linearized plasmids. Traditional integrative vectors contain the wildtype sequence of a prototrophic selection marker, such as *leu1+* or *ura4+* in case of the widely used pJK148 and pJK210 vectors, respectively. Upon linearization, plasmids can restore prototrophy upon recombination into corresponding genomic loci, which in the parental genome carry inactivating point mutations leading to auxotrophy, such as *leu1-32* or *ura4-294* (Keeney and Boeke, 1994). While efficient and site-specific, these recombination events also lead to up to 20% of transformants with multiple integration events (two or more copies integrated in tandem in the genome Keeney and Boeke, 1994). Even upon integration of the plasmid in single copy, the procedure leads to duplication of the auxotrophic marker genomic sequence (Fig.1A), which can cause locus instability. Indeed, we observed that strains in which fluorescent biosensors were introduced on such integrative vectors occasionally gave rise to cells lacking the probe (data not shown). While mutations in the gene encoding the probe could lead to apparent loss of signal, spontaneous mutations in wildtype fission yeast are rare (2×10^−10^ for base substitutions Farlow et al., 2015) and thus unlikely to account for the recurrent events we observed with different probes and strains. Instead, we speculated that the probe was lost because of recombination between the genomic repeats created by integrative vectors (Fig. 1A).

**Figure 1.**
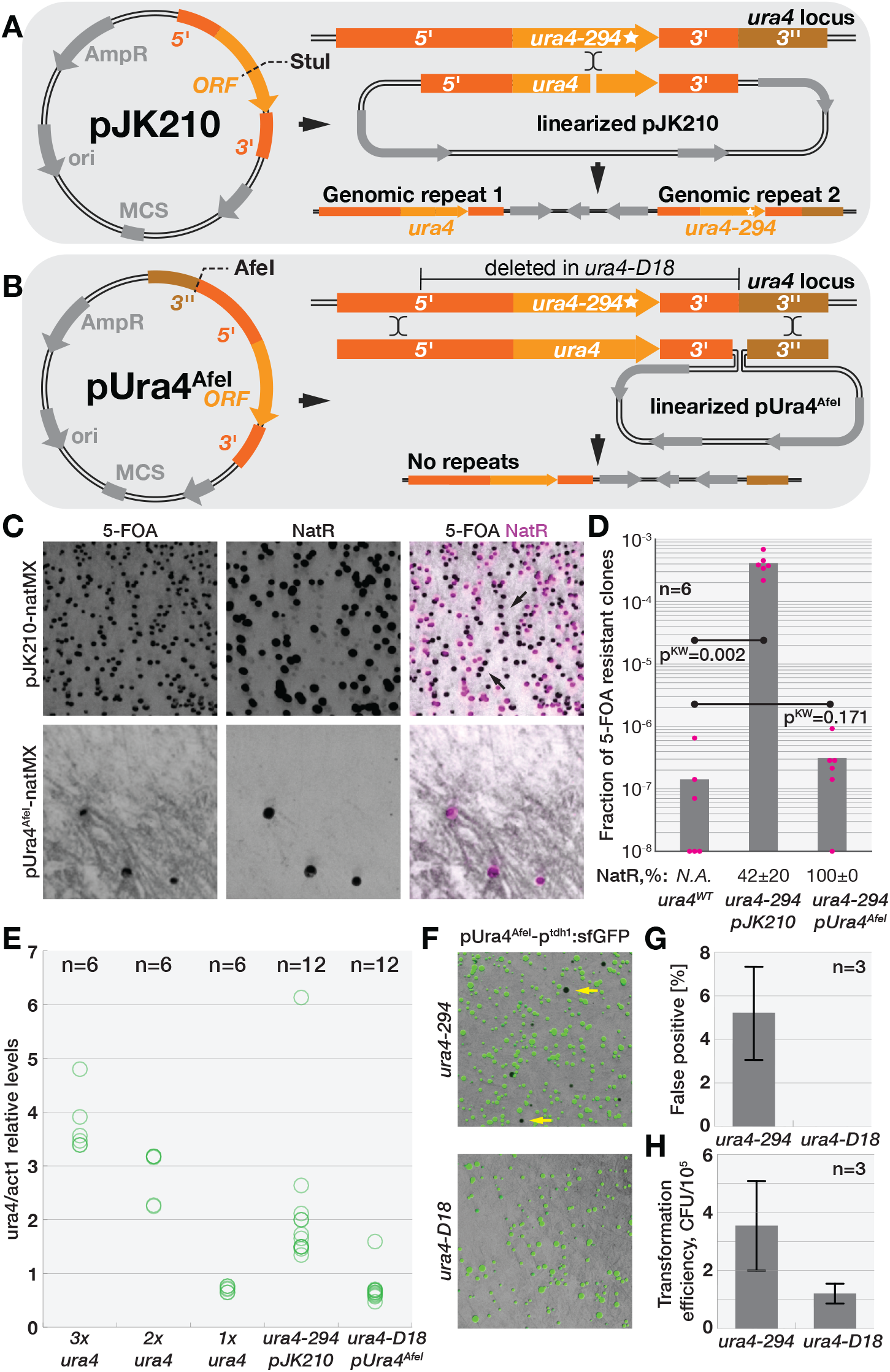
Two homology arms promote stable vector integration into the chromosome. **(A)** Schematic of the pJK210 vector (left panel) and its integration into the genomic *ura4-294* locus post-linearization (right panel). 5′, ORF and 3′ refer to *ura4* gene sequences as indicated in the genomic locus. AmpR encodes for ampicillin resistance and ori stands for the ColE1 bacterial replication origin. The unlabelled arrow represents an F1 replication origin. A single site crossover leads to the integration of the vector and duplication of the target locus (for detailed explanation see Jasin and Rothstein, 2013; Lee et al., 2014). (B) Schematic of the pUra4^AfeI^ vector (left panel) and its integration into the genomic *ura4-294* locus post-linearization (right panel). 3” refers to the indicated sequences downstream of the 3’ segment. Note that the two homology regions are separated by the *AfeI* linearization site. Crossovers at both homology arms integrate the vector into the genome without duplication of the target locus. Note that the *ura4-294* point mutation in the ORF can be rescued by a single crossover that does not lead to vector integration. The *ura4-D18* mutant, which lacks the indicated fragment, only has homology with the distal ends of the homology arms, ensuring that the vector integrates along with the *ura4+* selection cassette. (C) Assessment of integrant stability. Yeast strains transformed with the plasmids pJK210 (top) or pUra4^AfeI^ (bottom) in which the *ura4+* and natMX6 selection cassettes were introduced were grown in non-selective conditions for three days and 1.4×10^7^ of cells plated on media with 5-FOA (left panels), which is lethal to cells encoding a functional *ura4+* gene. After colonies developed, we replica plated them onto media containing nourseothricin (middle panels). False-colored images from the two plates are overlaid on the right panel. Note that numerous colonies develop if selection markers were originally introduced using pJK210 but not pUra4^AfeI^ vector. Approximately half of the 5-FOA-resistant clones from the pJK210 integration also lost the natMX6 cassette, which was maintained in all clones obtained from pUra4^AfeI^-transformed strain. **(D)** The graph quantifies the fraction of 5-FOA resistant clones upon plating 1.4×10^7^ cells. The percentage of nourseothricin resistance is shown at the bottom of the graph. **(E)** Quantitative PCR results comparing the relative abundance of *act1* and *ura4* genomic loci for strains containing the indicated number of *ura4* loci, and for 12 strains obtained by transformation with indicated plasmids each. **(F)** Measure of false-positive integrations. Indicated *ura4* mutant alleles were transformed with a cassette for sfGFP expression cloned into the pUra4^AfeI^ vector and selected for growth on media lacking uracil. The colonies that developed were imaged in white light (grayscale image) and green fluorescent channel (green) and the images were overlaid. The arrows point to false positive colonies that lack the fluorescent marker but contain the functional *ura4* cassette. **(G)** Quantification of percentages of false positive colonies observed in (F). **(H)** Quantification of overall transformation efficiencies observed in (F).

To quantify the instability of pJK210 integrants, we introduced a nourseothricin resistance cassette (natMX) into the pJK210 vector, which carries a *ura4+* gene as a selection marker. We transformed auxotrophic cells carrying the point mutation *ura4-294* and selected for uracil prototrophs with nourseothricin resistance. We subsequently cultured these transformants in non-selective media for three days and plated 1.4×10^7^ cells onto media containing 5-fluoroorotic acid (5-FOA), which acts as a counter selection against the *ura4+* gene (Fig. 1C). Inactivation of *ura4* in pJK210 integration strains occurred at a frequency that was three orders of magnitude above the rates we observed for a wildtype prototroph strain (Fig. 1D, 4.1± 1.6×10^−4^ for pJK210 transformants and 1.4±2.5×10^−7^ for wildtype cells, N=6). Furthermore, 58.0+20.3% of transformants with inactivated *ura4* gene also lost the resistance to nourseothricin, suggesting loss of vector sequences. Our results suggest that recombination between the repeats introduced upon pJK210 integration are a likely cause for the instability of the integrated construct.

We aimed to create a stable integration vector by developing a new pUra4^AfeI^ vector (Fig. 1B), which integrates at the *ura4* locus without creating genomic repeats. pUra4^AfeI^ linearization at the *AfeI* site produces two separate homology regions (Fig. 1B): The first homology arm contains the functional *ura4* cassette (5’ region, ORF, 3’ region). The second arm is homologous to sequences just downstream of the *ura4* cassette (3’’ region). Integration of the linearized fragment relies on recombination at both homology regions, which replaces the genomic *ura4* sequences and thus avoids repeat formation (Fig. 1B). Integration is confirmed using three diagnostic PCRs, which probe for correct integration on both sides of the linearized vector and for an increase in the distance between the 3’ and 3’’ *ura4* genomic regions (Fig. S1; see Materials and Methods for details). We introduced the natMX6 cassette using the pUra4^AfeI^ vector into the *ura4-294* mutant strain. Nourseothricin-resistant, uracil prototroph transformants exhibited a stable *ura4* locus with a frequency of cells resistant to 5-FOA similar to that of wildtype prototrophs (Fig. 1D). Importantly, all the 5-FOA-resistant clones that arose in the population maintained the nourseothricin resistance, indicating continued presence of vector sequence (Fig. 1C). We conclude that exogenous DNA can be stably introduced in the genome at a defined locus by avoiding the formation of genomic repeats.

To monitor the presence or absence of genomic repeats upon vector integration, we used quantitative PCR (qPCR). We first characterized the sensitivity of the assay by examining strains carrying the *ura4* gene in single (wildtype *ura4+*), two (*pak2∆::ura4+ ura4+*) and three (*myo51∆::ura4+ pak2∆::ura4+ ura4+*) copies. The qPCR results clearly reflected an increase in *ura4* copy number as compared to the *act1* gene, which we used as a reference locus (Fig. 1E). We then assayed the genomic DNA of twelve clones obtained by transforming pJK210 and pUra4^AfeI^ vectors. The majority of pJK210 transformants showed two copies of the *ura4* gene, consistent with duplication of the *ura4* locus upon vector integration. In addition, one clone showed a >3x *ura4* signal, revealing a multiple integration event (Fig. 1E), consistent with previous data that pJK210 can integrate in multiple tandem copies (Keeney and Boeke, 1994). In the case of pUra4^AfeI^, all but one clone showed *ura4* present in single copy, with one clone possibly present in two copies (Fig. 1E). Thus, pUra4^AfeI^ integrates in the genome without causing duplication. In addition, these results show that the majority of transformants result in single-copy integrations, consistent with the vector design.

To quantify the rates of false positive clones produced with the pUra4^AfeI^ vector (*e.g.* clones that are uracil prototroph but have not integrated the vector), we used pUra4^AfeI^ to introduce a locus encoding high levels of sfGFP into *ura4-294* cells. Blue light illumination showed that transformant colonies obtained after uracil selection were green due to sfGFP expression. However, sfGFP expression was absent from 5.2±2.1% of transformants (Fig. 1F, top panel; 1G). False positive transformants may arise from a double crossover that only spans the *ura4-294* point-mutation and restores wildtype *ura4+* gene without integrating any vector-specific sequences. The commonly used *ura4-D18* mutant allele (Grimm et al., 1988) has the entire open reading frame of *ura4* gene deleted, leaving homology only to the tips of homology arms in the linearized pUra4^AfeI^ (Fig. 1B). Transformation of pUra4^AfeI^ into cells harboring *ura4-D18* resulted in false positive rates below the detection limit of our assay (~ 0.2%; Fig. 1F, bottom panel; 1G). While the routine transformation protocol still yielded hundreds of transformants, the transformation efficiency of pUra4^AfeI^ into the *ura4-D18* strains decreased three-fold as compared to *ura4-294* (Fig. 1H, 1.2±0.3×10^−5^ and 3.5±1.5×10^−5^ for the *ura4* deletion and point mutant respectively). We conclude that restricting homology with the genome to the edges of the linearized vector leads to decreased rates of false positive transformants.

Taken together, we show that the pUra4^AfeI^ vector can be used to reliably integrate sequences in single copy into the genome without producing genomic repeats. We find that minimal rates of false positive transformants are achieved when targeting the *ura4-D18* deletion mutant locus.

### A series of stable integration vectors

Experiments increasingly rely on simultaneously monitoring multiple probes and markers. We thus developed a series of vectors similar to pUra4^AfeI^ but targeting the additional *ade6*, *lys3* and *his5* loci. For each locus, we cloned the plasmid backbone with one homology arm that contains the 5’ region, ORF and 3’ region, which is preceded by a linearization site and a second homology arm targeting the downstream 3’’ region (Fig. 2A). We named these plasmids pAde6^PmeI^, pLys3^BstZ17I^ and pHis5^StuI^.

**Figure 2.**
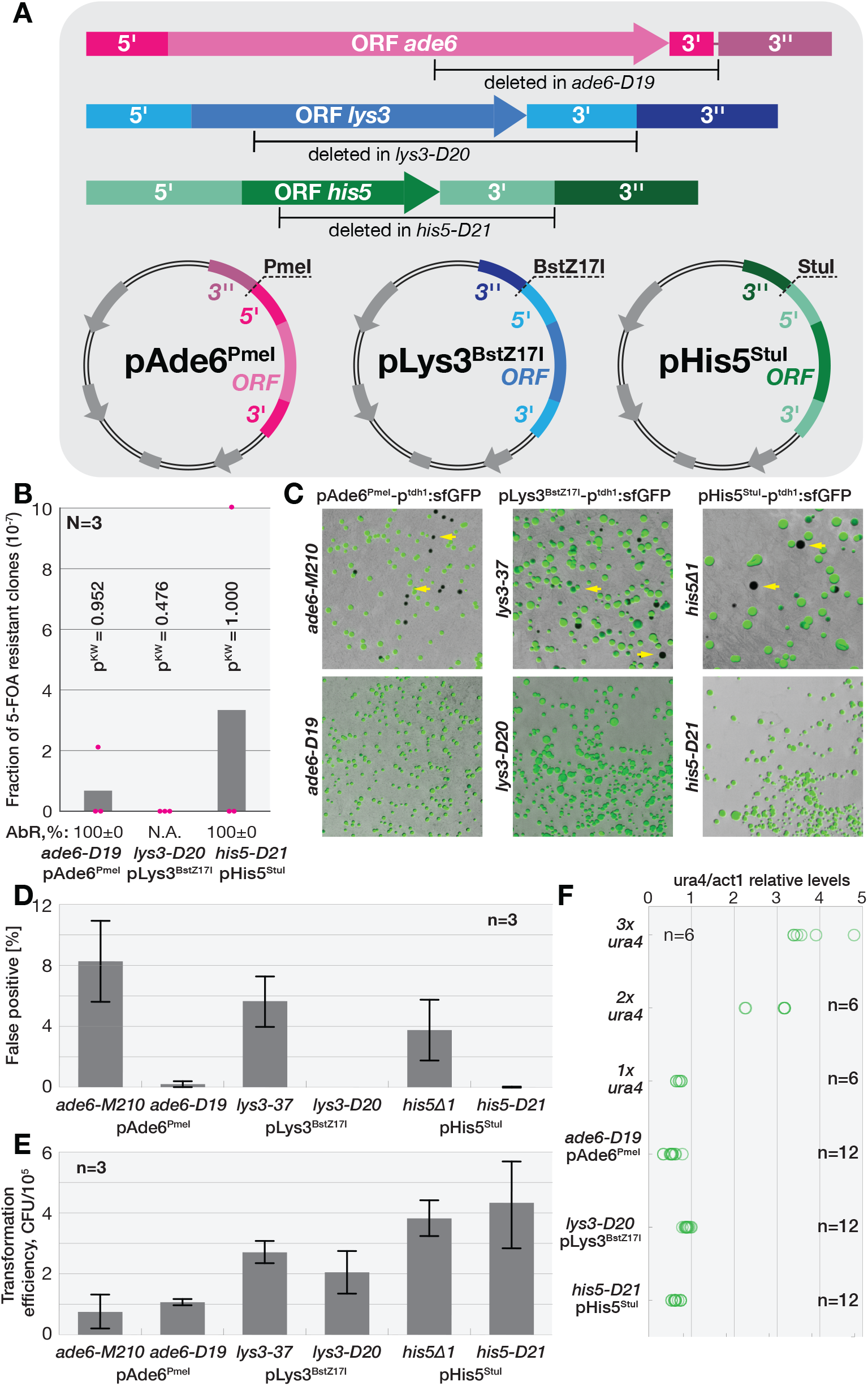
Vector series for stable chromosomal integration. **(A)** Schematics of the genomic loci (top) targeted by pAde6^PmeI^, pLys3^Bstz17I^ and pHis5^StuI^ vectors (bottom). The segments that are deleted in the mutants *ade6-D19*, *lys3-D21* and *his5-D21* are also indicated. Notations are as in Fig 1B.**(B)** Assessment of integrant stability. Yeast strains, where the *ura4* and antibiotic selection cassettes were introduced on indicated plasmids, were grown in non-selective conditions for three days and plated onto media with 5-FOA. After colonies developed, we replica plated them onto media containing the antibiotic. The graph quantifies the rates of 5-FOA resistant clones that formed upon plating 1.4×10^7^ cells. The percentage of antibiotic-resistant colonies is indicated at the bottom. **(C)** Measure of false-positive integrations. Indicated mutant alleles were transformed with a cassette for sfGFP expression cloned into indicated vectors and prototrophs were selected. The colonies that developed were imaged in white light (grayscale image) and green fluorescent channel (green) and the images were overlaid. The arrows point to false positive prototrophic colonies that lack the fluorescent marker. **(D)** Quantification of percentage of false positive colonies observed in (C). **(E)** Quantification of overall transformation efficiencies observed in (C). **(F)** Quantitative PCR results comparing relative abundance of *act1* and *ura4* genomic loci for strains containing indicated number of *ura4* loci (same data as in Fig. 1E) and for 12 strains obtained by transformation with indicated plasmids that carry the *ura4* gene. Note that all transformants exhibit *ura4/act1* relative levels indicative of a single integration event.

We first tested the stability of pAde6^PmeI^, pLys3^BstZ17I^ and pHis5^StuI^ integrants. To this aim, we simultaneously introduced both *ura4+* and antibiotic resistance cassettes into the vectors, linearized them and transformed them into *ura4-D18* uracil auxotroph yeast cells. Note that these strains were prototroph for adenine, lysine and histidine. We selected clones that were both uracil prototroph and antibiotic resistant. We then monitored the stability of the integrated locus by counter-selecting against *ura4+* using 5-FOA. The inactivation of the *ura4* gene occurred at frequencies similar to wildtype cells (Fig. 2B, 7.1±12.4×10^−8^ for *ade6* locus, 3.3±5.8×10^−8^ for *his5* locus, and bellow detection limit for the *lys3* locus), suggesting it was due to spontaneous mutations. Importantly, the strains with inactivated *ura4* gene invariably maintained the antibiotic resistance (Fig. 2B). We concluded that, as with pUra4^AfeI^, transformation with pAde6^PmeI^, pLys3^BstZ17I^ and pHis5^StuI^ produces stable integrants. We note that we also attempted the same strategy to target the *leu1* locus with pLeu1^StuI^ but found that the selective marker was rapidly lost in most transformants after removing the selective pressure. While we did not pursue this issue further, a possible explanation is that the *leu1* genomic region contains a replication origin, which allows the plasmid to exist as an unstable, episomal element.

We proceeded to assess the rate of false positive transformants with the newly-designed vectors. To this aim, we introduced a cassette that expresses high levels of sfGFP in pAde6^PmeI^, pLys3^BstZ17I^ and pHis5^StuI^ and integrated the vectors into the genome of *ade6-M210*, *lys3-37* or *his5∆1* auxotrophic strains, respectively. After selection for prototrophs we could distinguish colonies that do and do not express the fluorophore, and thus determine the proportion of false positives (Fig. 2C-D; 8.2±2.6% of false positives for *ade6-M210*, 5.6±1.7% for *lys3-37* and 3.8±2.0% for *his5∆1* alleles). We suspected that we could decrease the false positive rates by designing new auxotrophic deletion alleles (detailed in Materials and Methods). Specifically, we engineered *ade6-D19*, *lys3-D20* and *his5-D21* mutants that lack most of the ORF and the immediate 3’ region, and thus must recombine at both upstream and downstream regions to introduce the selection marker into the genome. Using these strains to integrate the fluorescent cassette into the genome almost completely abolished appearance of false positive clones (Fig. 2C-D; 0.21±0.19% of false positives for *ade6-D19*, below detection limit for *lys3-D20* and 6.5±11.3 x10^−3^ % for *his5-D21* alleles). Using our new deletion alleles had little effect on overall transformation efficiencies (Fig. 2E). Finally, we used our plasmids to integrate the *ura4+* cassette at either *ade6*, *lys3* or *his5* locus of the *ura4-D18* mutant. This allowed us to then monitor the number of plasmid integrants by the qPCR assay described above, which compared relative levels of *ura4* and *act1* sequences in the genome (Fig. 2F). We tested 12 transformants with each plasmid and found that all underwent a single integration event.

Taken together, our results indicate that the pAde6^PmeI^, pLys3^BstZ17I^ and pHis5^StuI^, as well as pUra4^AfeI^, can be used to introduce desired sequences into the fission yeast genome almost without any copy number variation. Furthermore, using auxotrophic deletion alleles with restricted regions of homology ensures almost negligible rates of false positive transformants.

### Expanding the usage of stable integration vectors

Using the above-described vectors requires target strains to be auxotroph for the required loci. To circumvent this requirement, we decided to introduce additional dominant selection markers. We cloned the antibiotic resistance cassettes kanMX6, natMX6, hphMX6, bleMX6 and bsdMX6 into our vectors (Bähler et al., 1998; Hentges et al., 2005; Kimura et al., 1994; Sato et al., 2005; Wach et al., 1994; Table S1 and Fig. 3A; details in Materials and Methods) in between two MCS (Multiple Cloning Sites) to allow for their easy exchange between vectors. Transformation of linearized vectors into wildtype strains followed by antibiotic selection readily produced the desired clones (data not shown). This set of vectors targeting 4 distinct genomic sites with 5 different dominant selection markers, currently composed of 5 distinct vectors (Table S1), can be used to clone any sequence of interest in either MCS and stably introduce it in the yeast genome in essentially any yeast strain.

**Figure 3.**
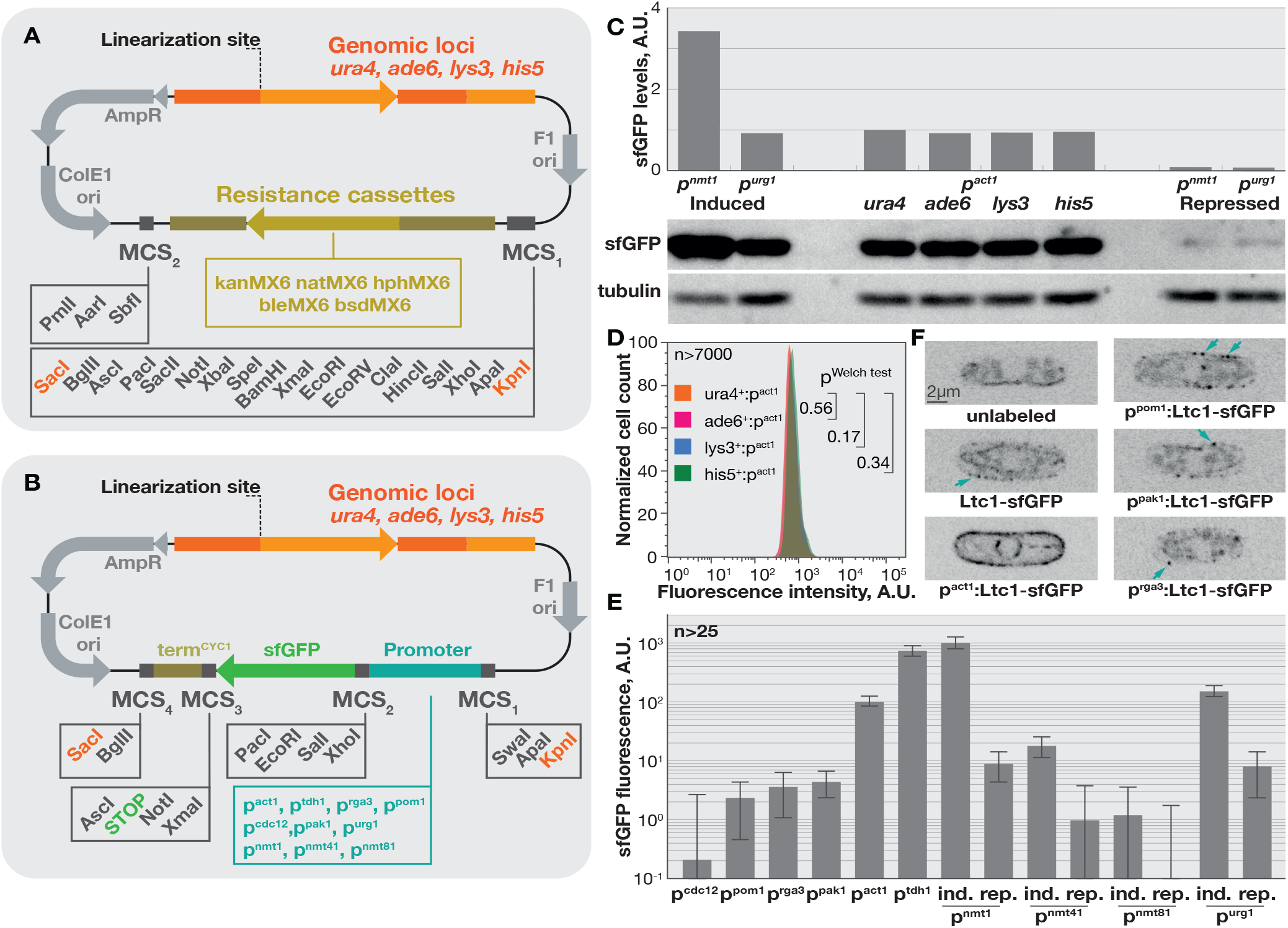
Expanding the Stable Integration Vector toolbox. **(A)** Schematics of cloned vectors with indicated genomic targeting loci (orange), antibiotic resistances (mustard) and multiple cloning sites (MCS). Note that not all sites are unique in all vectors. Please check in the vector sequences provided in supplement before planning your cloning. **(B)** Schematics of cloned vectors with indicated genomic targeting loci (orange), promoters (turquoise) and multiple cloning sites (MCS). The STOP codon is placed between restriction enzyme sites which allows to use the vectors for both N- and C-terminal sfGFP tagging. Note that not all sites are unique in all vectors, and that *KpnI* and *SacI* (highlighted) can be used to shuttle the entire constructs between vectors of this and the series presented in (A). **(C)** Quantification (top panel) of western blots against sfGFP (middle panel) and tubulin (bottom panel, loading control) performed on lysates obtained from cells where sfGFP was expressed from the indicated promoters. **(D)** Fluorescence profiles obtained by flow cytometry from cells with indicated genotypes. Note that there are no significant differences in expression of sfGFP from the *act1* promoter integrated at any of the four different genomic loci as suggested by the Welch tests p-values that are also reported. **(E)** Quantification of fluorescence emitted by the sfGFP that was expressed from the indicated constitutive, induced and repressed promoters. Mean values and standard deviation are reported. **(F)** Micrographs of Ltc1-sfGFP expressed from indicated promoters of different strengths and imaged with indicated exposure times. Please note that fluorescent foci (arrows) are barely notable above cell autofluorescence when Ltc1-sfGFP is expressed from the native *ltc1* locus but become clearly evident when using the *pom1*, *rga3*, or *pak1* promoters. Also note that using the strong *act1* promoter results in ectopic localization throughout the endoplasmic reticulum.

In most experimental settings, control over gene expression levels is desirable. To this aim, we constructed a second set of vectors, in which 10 different promoters drive the expression of sfGFP followed by the transcriptional terminator of the budding yeast *CYC1* gene. These include the inducible promoters *p^nmt1^*, *p^nmt41^*, *p^nmt81^* and *p^urg1^* and six distinct constitutive promoters active during mitotic growth. Multiple cloning sites are present in between each of the fragments to allow their easy exchange, as well as to use the plasmid for both C- and N-terminal fusion proteins with sfGFP (Fig. 3B; Note that the STOP codon is within the MCS_3_). Next, we compared the induction strength of the 10 promoters active during mitotic growth. First, we obtained whole cell lysates from selected strains and analyzed sfGFP levels by western blotting using tubulin as a loading control. Both *nmt1* and *urg1* inducible promoters showed robust fold-differences between non-induced and induced conditions, with the *nmt1* promoter expressing at higher levels (Fig. 3C). Importantly, we found that the constitutive *act1* promoter, which leads to similar expression levels as the *urg1* promoter in presence of uracil, drives similar expression levels when integrated at any of the four genomic loci (Fig. 3C). Comparison of sfGFP expression from the *act1* promoter integrated at different genomic loci by flow cytometry confirmed that the integration site does not influence expression levels (Fig. 3D). This suggests that there are no strong differences in chromatin accessibility for transcription between these four genomic loci, as observed for other loci (Allshire and Ekwall, 2015), at least in otherwise wildtype cells.

To compare the activity of other promoters driving sfGFP expression, we quantified the fluorescence detected by spinning-disk confocal microscopy (Fig. 3E; see Materials and Methods for details). As observed by western blotting, the inducible promoters showed robust expression increase in induced conditions, with the *nmt1* promoter yielding the highest expression level, followed by *p^urg1^*, *p^nmt41^* and *p^nmt81^*. Note however that *p^urg1^* was expressed only about 20-fold upon uracil addition, as previously reported (Watson et al., 2013). The *tdh1* promoter was the highest-expressing constitutive promoter, yielding sfGFP expression levels just slightly lower than the fully-induced *nmt1* promoter. This is thus a good promoter for very strong constitutive expression. The *act1* promoter led to about 7-fold lower expression levels, similar to induced *urg1*. The *pak1*, *rga3* and *pom1* promoters exhibited a further 23-, 27- and 42-fold decrease in levels, respectively. The *cdc12* promoter was about 10-fold weaker, yielding barely detectable GFP levels. We were unable to reliably detect cytosolic sfGFP when expressed from the repressed *nmt81* promoter. However, this is likely due to limitations of the assay in detecting a weak cytosolic signal against background organellar fluorescence because we obtained evidence that *cdc12* and *nmt81* promoters in the off-state lead to detectable biological activity (Hachet et al, 2011; data not shown). We further used the *act1, pak1, rga3* and *pom1* promoters, all of which lead to detectable levels of cytosolic GFP, to express Ltc1, a protein localized at ER-PM contact sites (Fig. 3F; Marek et al., 2019). The localization of Ltc1, which was difficult to detect from its native promoter, showed a more prominent localization pattern when mildly overexpressed from *pak1*, *pom1* or *rga3* promoters. Stronger overexpression from the *act1* promoter resulted in ectopic localization throughout the ER. This example illustrates how the range of promoters driving expression over three orders of magnitude will allow tweaking of gene expression levels to reveal biological insight.

In summary, we provide three sets of stable, single-integration vectors: 1) pUra4^AfeI^, pAde6^PmeI^, pLys3^BstZ17I^ and pHis5^StuI^, which lead to stable, single-copy integrations when transformed into strains auxotroph for the corresponding marker gene (Fig. 1 and 2); 2) a derived set containing additional antibiotic resistance cassettes, which allow their integration independently of the host strain genotype also in prototrophic strains (Fig. 3A); and 3) a set of vectors with 10 distinct promoters to drive gene expression at defined levels (Fig. 3B). This third set also needs to be transformed into auxotrophic strains. However, we note that subcloning with *KpnI* and *SacI* restriction enzymes allows introducing the DNA fragment containing promoter, sfGFP and MCSs from the third plasmid set into the second set of vectors next to the antibiotic resistance cassette (see Box 1) for guidelines and further examples of modularity). This exemplifies the modularity of the set of plasmids we created and their possible expansion.

### Using single integration vectors to express live cell biology probes

Synthetic probes are routinely used in cell biology to monitor molecular dynamics and activity. For reliable quantifications and phenotype comparisons it is imperative that probe levels be comparable between samples. Because our single-integration vectors show invariant copy number, we used them to introduce a number of fluorescent markers into cells (Fig. 4). We used different loci and antibiotics, which allows to quickly combine multiple probes in the same cell by genetic crosses. Furthermore, we used three distinct fluorescent tags: sfGFP (Pédelacq et al., 2006), mCherry (Snaith et al., 2005) and the blue fluorophore mTagBFP2 (Subach et al., 2011), which produces a signal that is efficiently separated from GFP and mCherry by standard DAPI filters (see Materials and Methods). We used strong promoters to drive expression of cytosolic blue, green and red fluorophore (Fig. 4A) which can be used to distinguish cells when simultaneously imaging multiple strains. We also placed the three fluorophores under mating type specific promoters that are active only in P-gametes (*p^map3^*) or M-gametes (*p^mam1*^*). This allowed us to clearly differentiate gametes during mating (Fig. 4B, Mov. S1). To visualize the plasma membrane, we fused the three fluorophores to an amphipathic helix from RitC (Fig. 4C). We targeted each fluorophore to the nucleus using a SV40 nuclear localization sequence at either one (Fig. 4D) or both ends of the protein (Fig. 4E). To monitor active export of proteins from the nucleus, we fused sfGFP with the nuclear export sequences from Mia1/Alp7 and Wis1 fission yeast proteins (Fig. 4F). To monitor nuclear envelope integrity, we expressed three copies of mTagBPF2 in tandem, whose size largely prevents its nuclear influx in wildtype cells (Fig. 4G). We note that the 3mTagBFP2 also occasionally made cytosolic foci, possibly due to self-oligomerization.

**Figure 4.**
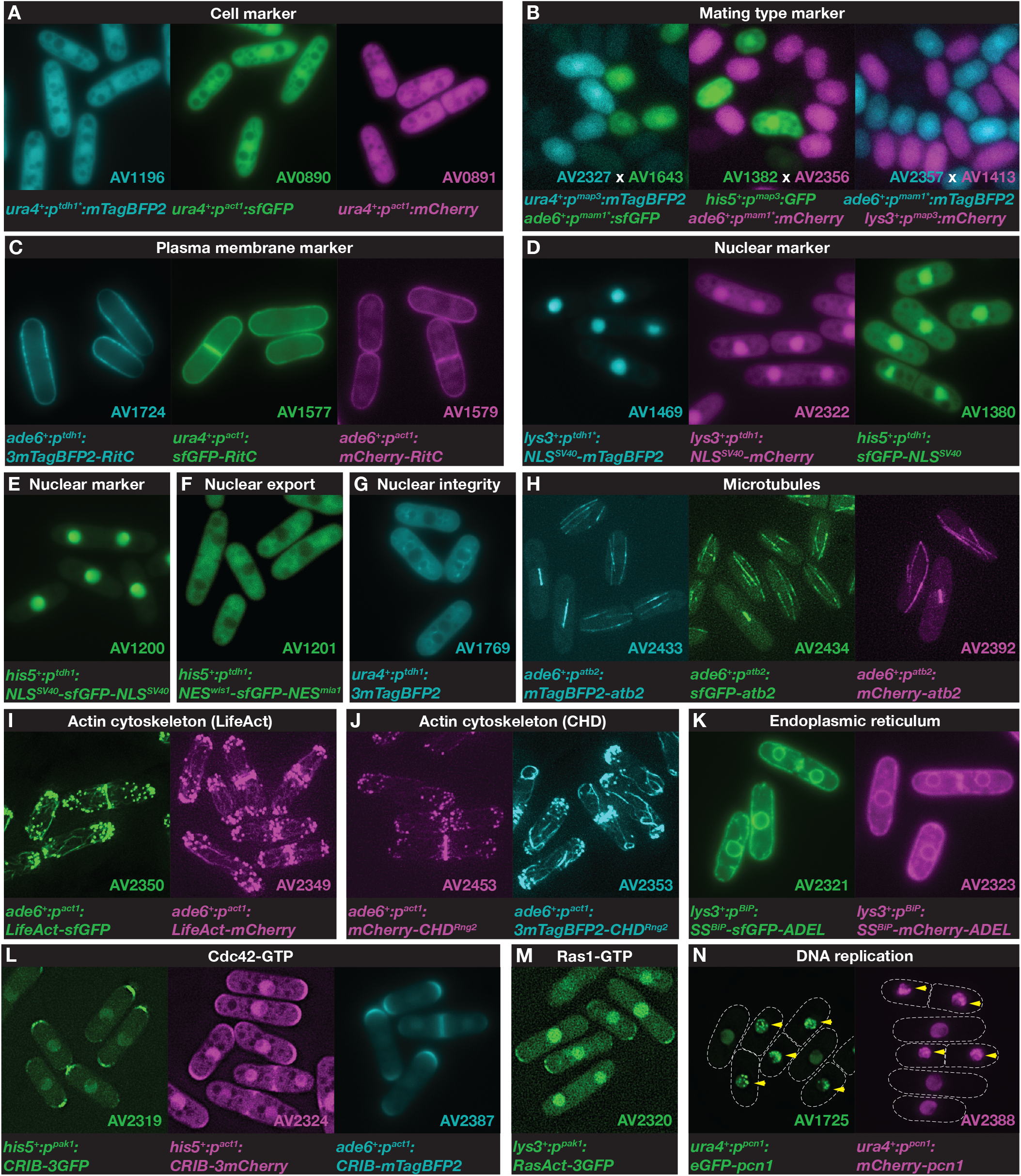
Panel of cell biology probes generated using SIV toolbox. Micrographs of cells expressing green, red and blue fluorescent probes to visualize **(A)** cells, **(B)** P- and M-gametes, **(C)** plasma membrane, **(D, E)** nucleus and/or nuclear import, **(F)** nuclear export, **(G)** nuclear integrity, **(H)** microtubules, **(I, J)** actin cytoskeleton, **(K)** endoplasmic reticulum, **(L)** active Cdc42, **(M)** active Ras1 and **(N)** DNA replication foci (arrowheads). Please see the figure and the text for detail of the constructs.

To monitor microtubules, we introduced the three fluorophores at the N-terminus of α-tubulin (*atb2*) and used its 5’ and 3’ regulatory sequences to drive expression of the construct as a second copy without perturbing the native α-tubulin locus (Fig. 4H, Mov. S1). We monitored the actin cytoskeleton using the LifeAct probe (Riedl et al., 2008) fused with a red or a green fluorophore (Fig. 4I, Mov. S1). We also expressed LifeAct fused to mTagBFP2. However, these cells exhibited growth defects (data not shown). As an alternative probe to visualize the actin cytoskeleton, we fused the calponin homology domain of Rng2 protein (Wang et al., 2004) with either red or blue fluorescent protein (Fig. 4J, Mov. S1). We note that cells expressing mTagBFP2-CHD exhibited abnormally abundant, likely stabilized actin cables. We fused the N-terminal signal sequence from BiP to target either sfGFP or mCherry to the endoplasmic reticulum and ensured its retention in the ER with the C-terminal ADEL sequence (Fig. 4K, Zhang et al., 2010). We also attempted to generate a blue ER marker but did not observe any fluorescence (data not shown). We expressed the CRIB-3GFP probe to monitor the GTP-bound form of the small GTPase Cdc42 using the *p^pak1^* promoter as previously reported (note that this CRIB probe is 27-aa shorter than that described in Tatebe et al., 2008; Fig. 4L, left panel, Mov. S1). CRIB fused to triple mCherry or mTagBFP2 was reliably observed only when the construct was expressed using the stronger *p^act1^* promoter (Fig. 4L, middle and right panels, Mov. S1). We used the RasAct probe to monitor the GTP-bound form of the small GTPase Ras1 as previously reported (Merlini et al., 2018; Fig. 4M, Mov. S1). Finally, to monitor cell cycle progression, green and red fluorescent proteins were fused with Pcn1, the PCNA component of the replisome, and the construct was expressed in addition to the native gene (Meister et al., 2007; Fig. 4N, Mov. S1). Pcn1 produced a uniform nuclear signal except in S-phase cells undergoing DNA replication when distinct fluorescent foci representing replication factories are observed (Fig. 4N, arrowheads).

Taken together our work provides a panel of frequently used live cell probes that are stably expressed from a single genomic copy introduced using the vectors we developed.

## Discussion

We present here a large series of modular vectors for stable, single-copy integration in the fission yeast genome. This vector series expands the genetic toolbox of the popular fission yeast model *Schizosaccharomyces pombe*. For the reasons highlighted below, these vector currently are the best tools to introduce foreign genetic material in the genome in a controlled manner. We thus encourage their wide use and further development. The vectors and strains developed herein are available from the Japanese National BioResource Project (NBRP; http://yeast.nig.ac.jp/yeast/) and the non-profit plasmid repository Addgene (https://www.addgene.org/).

### Stable and single-copy genomic integration

The controlled expression of desired genetic information is a powerful way to probe gene functionality. The popular model *Schizosaccharomyces pombe* can easily be transformed with plasmids, which have been the vector of choice for introduction of exogenous DNA. However, the fact that fission yeast centromeres are large, complex genomic elements has hindered development of centromeric plasmids such as those developed in budding yeast (Clarke and Carbon, 1980). Furthermore, fission yeast cells do not naturally carry plasmids such as the budding yeast 2µ plasmid, which carries a partitioning system (Chan et al., 2013; Strope et al., 2015). Instead, fission yeast research has relied on either autonomously replicating episomal plasmids or on vectors integrating into the genome. The episomal vectors, such as the widely used pREP series (Maundrell, 1993), segregate randomly between daughter cells at division, which results in a copy number variation within the clonal population. Furthermore, the plasmid selection has to be continuously applied or the plasmid is rapidly lost, in particular as cells go through sexual reproduction. Most integrative plasmids (Keeney and Boeke, 1994; Maundrell, 1993) carry a single homology region that targets them into the desired locus of the genome. While efficient and precise, vector integration produces genomic repeats, which can recombine and remove the integrated segment. With a rate of plasmid sequence loss from integrant strains at ~5 × 10^−4^ over three days (Fig. 1D), integrated DNA that decreases cell fitness would be rapidly eliminated from the population. This is likely to be particularly prominent in genetic crosses, when genomic repeats misalign between parental chromosomes during meiotic recombination (Smith, 1976). In summary, most traditionally-used vectors suffer from instability and copy-number variation.

To overcome this problem, we developed pUra4^AfeI^, pAde6^PmeI^, pLys3^BstZ17I^ and pHis5^StuI^ plasmids, which rely on two homology arms to integrate into the genome (Fig. 1A and 2A). These vectors recombine specifically with their target genomic locus (Fig. S1B). Importantly, we have shown that integration of these vectors occurs without producing genomic repeats, and leads to a stable genomic copy that is not lost when grown without selection (Fig. 1C-D and 2B). The vectors also predominantly integrate as a single copy (Fig. 2F), which ensures that copy-number is not a confounding factor when comparing the biological activity of different constructs. Thus, these vectors allow reliable stable, single-copy genomic integration.

### Comparison with other vector series

Integrative plasmids that rely on two homology arms have previously been developed by the Sato and Hagan groups (Fennessy et al., 2014; Kakui et al., 2015). Whether these plasmids, which target different genomic loci than the ones used here, exhibit similar properties remains to be seen, as neither stability of integration nor number of integration events have been reported. It may be unwise to assume that all sites of integration behave similarly. Indeed, we found here that targeting the *leu1* locus with a plasmid carrying two homology arms (pLeu1^StuI^) did not systematically lead to stable genomic integration. One possible explanation is that the *leu1* sequence contains a replication origin, which may allow the cells to transiently maintain the plasmid as an episomal element if it is re-circularized by non-homologous end joining. Indeed, a potential replication origin resides at the *leu1* locus (originID: II-1983; Siow et al., 2012). The design of these previously-described plasmids generates another major difference with the set of integrative plasmids described here: their integration leads to disruption of the target genomic site, and thus the creation of auxotrophic strains, whereas integration of the plasmids presented here restores (or preserves) the target locus and thus forms prototrophic transformants. This may be beneficial for the study of many physiological pathways.

### Plasmid modularity and versatility

Because every experimental design is different and there is no single vector that can fit all needs, the vector series presented here has been developed with modularity in mind. Simple subcloning can easily re-target a construct to a different genomic locus, introduce a different selection marker, exchange the fluorescent tag or alter the level of expression. We already generated a set of highly used cell biology markers that can be combined through genetic crosses and are predominantly away from the most frequently targeted *ura4* locus. These can also be targeted to different loci or in other color, at will. We encourage the further expansion of the SIV series through introduction for instance of additional tags or inducible promoters (Kjaerulff and Nielsen, 2015; Ohira et al., 2017; Zilio et al., 2012).

The design choice will be dictated by the experiment (see Box 1 for recommendation on experimental design and plasmid use). For instance, while selecting for prototrophs is more cost effective, this requires transformation into an auxotrophic strain. For this, both strains carrying point mutations, which are present in most strain collections, and the new deletion alleles we constructed (*ade6-D19*, *lys3-D20* and *his5-D21*; Fig. 1F-G, 2C-D), as well as the previously-described *ura4-D18* allele (Grimm et al., 1988), are suitable. However, we find that prototrophic restoration of the selective marker without integration of the accompanying vector sequences occurs in up to 8% of the total transformants when using point mutants (Fig. 1F-G, 2C-D). Thus, strains with deletion alleles should be used when it is important to avoid false-positive transformants. For example, high rates of false-positive transformants may make it complicated to quantify how introducing genes of interest affects the capacity of cells to form colonies (Li and McLeod, 1996).

##### Box 1. Recommendation for use of the vectors

For best use of the tools described here, we make a few recommendations as follows:

1. Clone your construct in your plasmid of choice according to experimental design, bearing in mind that the restriction enzyme used for linearization (*AfeI*, *PmeI*, *BstZ17I* or *StuI* in pUra4^AfeI^, pAde6^PmeI^, pLys3^BstZ17I^ and pHis5^StuI^, respectively) should remain unique after cloning.

- In some cases, combinations of other restriction enzymes can be used for linearization as long as they produce a single linear fragment with two large homology arms (*e.g*. *RsrII* and *BlpI* can be used together on pAde6^PmeI^);
- Bear in mind that the multiple cloning sites between plasmids of the same sets are identical but not all restriction enzymes are unique in all cases. While plasmids carrying the antibiotic-resistance markers are designed to receive constructs from other plasmids by *SacI-KpnI* digestion, antibiotic markers can also be shuttled into the promoter series by using a unique site in either the *ColE1* or *AmpR* genes and *SacI* or *BglII*;
- Blunt restriction enzyme sites (*e.g*. *PmlI*, *SmaI*, *SwaI*) have been introduced in some of the plasmids to increase their modularity. Similarly, restriction sites that use compatible cohesive ends (*e.g*. *XhoI-SalI* pair) increase the ability to shift constructs between plasmids;
- In our experience, a single ligation reaction can readily combine three elements into a single plasmid (*e.g*. *p*^*act1*^ cut with *KpnI* and *EcoRI* from pAV0714, mTagBFP2 cut with *EcoRI* and *AscI* from pAV0471 and the vector sequences from pAV0783 cut with *KpnI* and *AscI*);
- Other protein tags can be cloned in place of the sfGFP sequence, preserving the STOP codon in the MCS after the tag, to allow for both N- and C-terminal tagging;
- In addition to the main 22 plasmids discussed in the text, we also provide 22 plasmids that were used in building the strains presented in Figure4. These include additional promoters (*e.g*. *the constitutive promoter p^atb2^* and *p*^*bip1*^, and the mating type-specific promoters *p*^*mam1\**^ and *p*^*map3*^), fluorophores (*e.g*. 3GFP, 3mCherry, 3mTagBFP2) and a terminator (*S. cerevisiae ADH1*) that may be better suited for some uses.
2. Linearize 700 μg of the plasmid and transform in the strain, using either prototrophy or antibiotic selection;
3. Verify correct integration by diagnostic PCRs on both sides of the integration sites with primers as described in Figure S1.

High rates of false-positive transformants may also increase the workload when screening transformants obtained using a plasmid library. Using deletion alleles in the parental strain precludes homologous recombination of the selective marker alone, but we note that unintended transformant genotypes may still rarely arise when integration is achieved by non-homologous repair (Chang et al., 2017; Fennessy et al., 2014). The vector variants containing an antibiotic selection marker (Fig. 3A) should also exhibit low false-positive rates when selecting for the antibiotic resistance but not when selecting for prototrophic transformants. The plethora of dominant antibiotic resistance markers is particularly useful since vectors can be directly integrated into most genetic backgrounds without the need for a pre-existing auxotrophy. The limit would only be reached in prototrophs already containing the 5 different antibiotic resistances used here.

The series of the promoters we provide induces expression levels that span more than three orders of magnitude and thus should accommodate for a wide range of applications. We note that, for the same promoter, we did not observe changes in expression levels between different genomic loci (Fig. 3C-D). However, we refrain from using different constructs to compare between conditions as genomic position effects may become evident only in certain backgrounds (Allshire and Ekwall, 2015). While we present the full vector series for the first time here, some plasmids have been used in published studies (Billault-Chaumartin and Martin, 2019; Gerganova et al., 2019; Lamas et al., 2019; Marek et al., 2019; Vještica et al., 2018) which illustrates their application in fission yeast research.

## Materials and methods

### Plasmids

All plasmids were generated using standard restriction enzyme cloning and InFusion system (TaKaRa Bio, Kusatsu, Japan). The list of all plasmids used in the study is available in Table S1. Sequences are available as annotated GenBank format files (see Supplemental Sequences) for all constructs used in the study. Plasmids that are of general use are available from Addgene (Watertown, USA; https://www.addgene.org) and the Japanese National BioResource Project (NBRP, Osaka, Japan; http://yeast.nig.ac.jp). All plasmids used to test the system, which are derivatives of the ones available from these resource centers, are available from the Martin lab upon reasonable request.

The plasmid backbones were amplified from pJK210 which itself was derived from pBluescript SK (+) and encode for ColE1 and F1 replication origins, ampicillin selection cassette, a multiple cloning site and T3, T7 and M13 primer sequences. Dominant selection markers kanMX6, natMX6, hphMX6, bleMX6 for yeast selection were amplified from the pFA6a vector series (Bähler et al., 1998; Hentges et al., 2005; Wach et al., 1994). The bsdMX6 selection marker against was amplified from the fission yeast strain VS6381, a kind gift from Viesturs Simanis (EPFL, Switzerland) and Masamitsu Sato (Waseda University, Japan). We noticed sequence variation in comparison to original sequences, but report that all markers were fully functional. All fission yeast sequences were amplified from the genomes of wildtype strains 968 (ySM1396), 972 (ySM995) or 975 (ySM1371).

For **promoter sequences** we used the sequences immediately upstream of the START codon. Promoter p^tdh1^ contains 1000bp, while p^tdh1*^ promoter contains 896bp (used only in pAV0471, pAV0765 pAV0532, pAV0569 - pAV0572). p^act1^ includes 822bp, p^rga3^ includes 1203bp, p^pom1^ includes 688bp, p^map3^ includes 2063bp, p^atb2^ includes 647bp, p^bip1^ includes 988bp, p^pcn1^ includes 1954bp, p^pak1^ includes 630bp, p^urg1^ includes 675bp and p^nmt1^ includes 1177bp. The p^nmt41^ and p^nmt81^ promoters were obtained by subsequently mutating the p^nmt1^ using nested PCR (with mutations as reported by Basi et al., 1993). p^mam1*^ includes 1751bp and was point mutated to remove the AfeI site.

For **terminator sequences** we used the sequences immediately after the STOP codon of *Saccharomyces cerevisiae* genes *ADH1* (229bp) and *CYC1* (250bp). We also used the 988bp downstream of the fission yeast *nmt1* gene STOP codon. In case of sequences designated as the *tdh1* terminator, we used a fragment containing the sequence between STOP+332bp and STOP+1032bp. We note that this fragment does not include the 3’UTR sequence of the *tdh1* gene but led to a notable increase in protein levels as compared to constructs lacking any terminator sequence.

Fluorescent protein sequences were obtained from lab stocks with the exception of **mTagBFP2** which was amplified from mTagBFP2-pBAD (Addgene, Watertown, USA; plasmid ID 54572). The 3mTagBFP2 was a kind gift from Serge Pelet (University of Lausanne, Switzerland)

All the probes are derived from previous publications. **Nuclear localization signal** uses the viral SV40 motif (PKKKRKV, Kalderon et al., 1984). **Nuclear export signals** were derived from fission yeast genes Mia1/Alp7 (EDLVIAMDQLNLEQ, Ling et al., 2009) and Wis1 (QPLSCSLRQLSISP, (Nguyen et al., 2002). The **CRIB probe** encodes amino acids 2-181 from *S. cerevisiae* protein Gic2 in contrast to CRIB construct used by (Tatebe et al., 2008) which used amino acids 1-208. The **RasAct probe** encodes Byr1 amino acids 65-180 in three tandem repeats (Merlini et al., 2018). The **LifeAct probe** encodes a MGVADLIKKFESISKE peptide (Riedl et al., 2008). The **CHD probe** contains amino acids 1-189 from the Rng2 protein (Wang et al., 2004). The N-terminal **signal sequence** comprising amino acids 1-25 from the BiP (Bip1) protein targets the fluorophore to the ER while the C-terminal ADEL motif ensures its retention in the ER (Zhang et al., 2010). **Microtubules** are visualized by full length Atb2 (Cassimeris and Tran, 2010) and **replication sites** by Pcn1 (Meister et al., 2007) both tagged at the N-terminus and expressed as a second copy.

### Growth conditions

Yeast cells were grown in standard fission yeast media at either 25°C or 30°C (Hagan, 2016) and using 200rpm rotators for liquid media. For genetic manipulations we used YES media. For selection we supplemented YES with 100 µg/ml G418/kanamycin (CatNo.G4185, Formedium, Norfolk, UK), 100 µg/ml nourseothricin (HKI, Jena, Germany), 50 µg/ml hygromycinB (CatNo.10687010 Invitrogen), 100 µg/ml zeocin (CatNo.R25001, ThermoFischer), and 15 µg/ml blasticidin-S (CatNo.R21001, ThermoFischer). 5-FOA (CatNo.003234, Fluorochem, Derbyshire, UK) was used at 1 mg/ml in EMM media supplemented with 50.25 µg/ml of uracil (U0750-100G, Sigma, St. Louis, USA). For western blotting, flow cytometry and imaging we used EMM media. Briefly, cells were precultured overnight in liquid EMM media and then again sub-cultured overnight to reach exponential phase. Repression of the *nmt* promoters was achieved by supplementing thiamine at final concentration of 5 µg/ml. Activation of the *nmt* promoters was done by washing out thiamine and growing cells for minimum 24 hours. *urg1* promoter was induced by addition of uracil at final concentration 250 µg/ml for at least 48 hrs. Conditions used to mate cells are detailed in (Vjestica et al., 2016). Briefly, cells were precultured overnight in MSL+N media and then sub-cultured overnight to reach exponential phase. Cells of opposite mating types were then mixed in equal amounts, washed three times and resuspended in MSL-N to final O.D._600nm_=1.5 and incubated at 30°C with 200 rpm agitation for 4-6 hours prior to mounting for imaging.

### Yeast strains

The list of all strains used in the study is available in Table S2. Strains of general use are available from the Japanese National BioResource Project (NBRP, Osaka, Japan; http://yeast.nig.ac.jp).

All fission yeast strains were obtained by standard lithium-acetate transformation protocol (Hagan, 2016). We used ~700ng of the linearized plasmid per transformation. Unless otherwise indicated, plasmids were linearized with a single restriction enzyme present in between the two homology regions (*e.g. AfeI* for pUra4^AfeI^). In some instances, subcloned constructs contained additional cutting site for the enzyme normally used to linearize the vector. As indicated in the yeast strain table (Table S2), we overcame this problem by using two enzymes that flank the regular linearization sites (*e.g. RsrII* and *BlpI* for pAV0782) which lead to slightly shorter homology arms. This did not affect the ability to obtain correct transformants.

The ***ura4-294***, ***ade6-M210***, ***lys3-37*** are commonly used *S. pombe* point mutant strains that were available in many lab stocks. The ***his5∆1*** strain was obtained from Dr Mohan Balasubramanian (Tang et al., 2011). *ade6-M210* carries a C1466T substitution, whereas the exact mutations of other alleles are not known. The auxotrophic ***ura4-D18*** mutation (Grimm et al., 1988), which lacks the 1.8kb HindIII fragment flanking the ura4 ORF, was obtained by crossing out from the Martin group yeast library (YSM1131). To generate auxotrophic ***ade6-D19**, **lys3-D20*** and ***his5-D21*** deletions, we first inserted the *ura4+* selection cassette into the pAde6^PmeI^, pLys3^BstZ17I^ and pHis5^StuI^ vector to obtain plasmids pAV0596, pAV0597 andpAV0598. The resulting plasmids were linearized to target their integration at the *ade6*, *lys3* and *his5* genomic loci, respectively. Each linearized plasmid was separately transformed into the yeast strain carrying the *ura4-D18* mutation. Selection for uracil prototrophy yielded strains where a functional *ura4+* cassette was now linked with either *ade6*, *lys3* or *his5* locus, while the native *ura4* locus carried the *ura4-D18* mutation. These strains were then transformed with DNA fragments spanning both sides of the *ura4+* integration site and carrying the *ade6-D19, lys3-D20* or *his5-D21* deletions (see Supplemental Sequences). Recombinants that remove the *ura4+* cassette and replace the locus with the deletion allele were selected on 5-FOA. The *ade6-D19* allele is lacking the fragment from STOP-776bp to STOP+159bp, the *lys3-D20* allele lacks sequences between STOP-895bp and STOP+367bp, and the *his5-D21* allele lacks the fragment from STOP-524bp to STOP+378bp. The correct transformants were confirmed by diagnostic PCRs (the PCR-P∆ indicated in the Fig. S1) and then backcrossed six times with the wildtype fission yeast strain 975 (YSM1371) to obtain the final strains. The *h90* strains were obtained through crosses with wildtype fission yeast strain 968 (YSM1396)

All other strains were obtained by transforming linearized plasmids into either auxotrophic strains and selecting for prototrophs, or into prototrophs and selecting for antibiotic resistance. Multiple transformants were genotyped to verify correct plasmid integration. Since expression levels between several transformants did not vary we assumed that they all carried a single integrated copy.

### gDNA extraction

gDNA was extracted using a protocol described by (Lõoke et al., 2011) with minor modifications. Briefly, approximately 5×10^7^ of freshly streaked cells was resuspend in 100 μl of the isolation buffer (250mM LiaAc, 1%SDS) in a microfuge tube. Samples were incubated for 10 minutes at 70°C and briefly vortexed. We added 300μl of 100% ethanol to each sample and briefly vortexed. Next, we spun down the samples at 15’000 g for 3 minutes. The pellet was washed with 70% ethanol twice. The pellet was briefly dried to remove traces of ethanol. We then dissolved the pellet in 100 μl of 5mM Tris-HCl and spun down the cell debris for 1 minute at 3’000g. The supernatant was transferred to a fresh tube and the concentration was adjusted to 100 µg/ml. The samples were used for both standard and quantitative PCR amplifications.

### Genotyping

Overview of the diagnostic PCR reactions used to confirm genomic integration of the vector at the desired locus is presented in Fig. S1. Sequences of the primers we used are available in Supplemental Table S3. Briefly, we tested recombination for each homology arm of the vectors by PCRs with one vector-specific and one genome-specific oligo (Fig. S1, right panel). Simultaneously we tested for the presence of the parental locus using two genome-specific oligos. We used different primers to test the prototrophic and auxotrophic parental loci. Only transformants where all three PCRs indicated correct integration were used further.

Standard PCRs were performed with 5µM primers, 5ng/µl of the gDNA and a polymerase made in-house using the following cycler program:

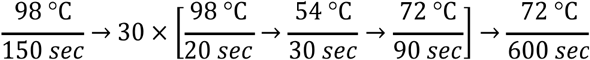

The *ade6* PCR-U (Fig S1) was done with commercial Kapa-Taq (Kapa Biosystems, Wilmington, USA) and the PCR program:

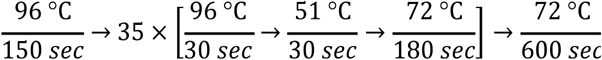

### Quantitative PCR

We used qPCR to monitor the number of plasmid integration events into the genome. The processed results are available in Supplemental Table S4 and raw EDS format data files in Source Data D1. Experiments were performed on the QuantStudio^TM^5 (Applied Biosystems, Foster City, USA) using SYBR Select Master Mix (Applied Biosystems, Foster City, USA; Cat No. 4472908) and primers at final concentration 0.2 µM. The final reaction volume was 10 µl. Each biological sample was evaluated by a technical triplicate. Technical outliers were rare and not removed. Each experiment included a negative control (water), which did not show any considerable signal amplification. An identical detection threshold of fluorescence intensity was set for all targets and all qPCR samples in the study and used to determine the cycle threshold (Ct) value. We monitored amplification of a *ura4* 142bp fragment using primers osm6182 (5’-GGCTGGGACAGCAATATCGT-3’) and osm6183 (5’-GCCTTCCAACCAGCTTCTCT-3’) and *act1* 149bp fragment using primers osm6178 (5’-GTGTTACCCACACTGTTCCCA-3’) and osm6179 (5’-TTCACGTTCGGCGGTAGTAG-3’). We first amplified the 10-fold dilution series of wildtype genomic DNA (extraction protocol is detailed above) at final concentrations in the range 0.01-100 ng/µl, which established primer efficiencies of 1.99 (R2 = 0.998) and 2.04 (R2 = 0.999) and intercepts of 18.53 and 19.06 for *ura4* and *act1* primer pairs, respectively. In subsequent experiments we used genomic DNA at final concentration 10 ng/µl and the formula

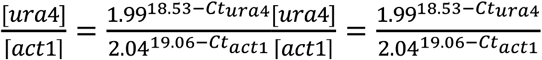

to compare relative levels of *ura4* and *act1* loci. We proceeded to show that we can reproducibly detect a relative increase in *ura4* locus copy number by using strains that carry one (ySM1371: *wt, ura4+*), two (AV0138: *pak2∆:ura4+ ura4+*) and three (AV0226: *myo52∆:ura4+ pak2∆:ura4+ ura4+*) copies of the *ura4* locus but only one *act1* locus. All subsequent experiments included each of these samples as positive controls.

To quantify the number of integrated copies of the vectors pJK210 and pUra4^AfeI^ we transformed the linearized vectors into *ura4-294* and *ura4-D18* auxotrophic strains, respectively, and extracted the genomic DNA from 12 prototrophic transformants. qPCR analysis of relative *ura4* and *act1* loci abundance was used to conclude the number of integration events. For vectors pAde6^PmeI^, pLys3^BstZ17I^ and pHis5^StuI^ we first introduced the *ura4* cassette to produce plasmids pAV0596, pAV0597 and pAV0597. Linearized vectors were transformed into *ade6-D19 ura4-D18, lys3-D20 ura4-D18* or *his5-D21 ura4-D18* double mutants, respectively, and selected for uracil prototrophy. 12 transformants from each transformation were then used to extract genomic DNA, establish relative levels of *act1* and *ura4* loci and conclude the number of integration events.

### Quantification of false positive integrants

To measure the proportion of false positive integrations we introduced sfGFP under the strong *tdh1* promoter into pUra4^AfeI^, pAde6^PmeI^, pLys3^BstZ17I^ and pHis5^StuI^ plasmids, which resulted in plasmids pAV0569, pAV0570, pAV0571 and pAV0572. After linearization, the plasmids were transformed into adequate auxotrophic strains and plated on media selecting for prototrophy. Once colonies formed, we imaged the selective plates using Fusion FX (Vilber, Collégien, France). While white light was used to observe all clones, we detect the green fluorescent protein using the 480nm LED illumination and the F535Y filter (535/50). This allowed us to distinguish between fluorescent and non-fluorescent clones. The percentage of non-fluorescent colonies was used as proportion of false-positive transformants. We report average value and standard deviation of three independent experiments.

### Quantification of stability of loci generated by plasmid integration

To quantify the stability of loci generated by integration of pJK210 and pUra4^AfeI^ plasmids, we exploited the 5-FOA counter-selection against *ura4+*. First, we cloned the natMX6 cassette into the two vectors to obtain plasmids pAV0584 and pAV0623. The resulting plasmids were linearized, transformed into cells carrying the *ura4-294* point mutation and for each transformation 6 nourseothricin-resistant uracil prototrophs were selected. To reveal any instability in the introduced loci, we grew the 12 transformant, and additional 4 clones of the wildtype prototrophic strain as a negative control, on non-selective (uracil supplemented, no nourseothricin) media for three days. The strains were first passaged twice on non-selective solid media over two days. We then precultured each clone in non-selective, liquid media overnight to exponential phase and plated 1.4×10^7^ cells onto 5-FOA media containing uracil, which selects for *ura4-* cells. 5-FOA resistant clones were replica plated them onto media containing nourseothricin to check whether the entire locus was lost. We report the average fraction of 5-FOA resistant clones out of 1.4×10^7^ cells that we plated, and the percentage of those that retained the nourseothricin resistance marker. Significance was determined using the Kruskal-Wallis test.

Since drugs to counter-select against *ade6+*, *lys3+* and *his5+* were not available, we introduced both the *ura4+* and an antibiotic cassette into pAde6^PmeI^, pLys3^BstZ17I^ and pHis5^StuI^ plasmids, which resulted in plasmids pAV0616, pAV0617, pAV0618. This allowed us to use the same strategy as above to quantify the stability of loci generated by our vector series, the only difference being that we performed it using 4 and not 6 initial transformant. Significance was determined using the Kruskal-Wallis test.

### Microscopy and image quantification

All images shown in Figure4 were obtained by wide-field microscopy performed on a DeltaVision platform (Applied Precision) composed of a customized inverted microscope (IX-71; Olympus), a UPlan Apochromat 100x/1.4 NA or 60x/1.4 NA oil objective, a camera (CoolSNAP HQ2; Photometrics), and a color combined unit illuminator (Insight SSI 7; Social Science Insights). Images were acquired using softWoRx v4.1.2 software (Applied Precision). The mTagBFP2 was imaged using the DAPI/FITC/TRITC/CY5^TM^ filterset which allows bandpass excitation (390/18) and emission (435/48). We used the GFP-mCherry^TM^ filterset to detect the green (Ex: 475/28, Em: 525/50) and red (Ex: 575/25; Em: 632/60) fluorescent proteins. We imaged a different number of Z-sections to best capture the structure of interest. We present either a single Z-plane or a projection image. All image processing was done using standard ImageJ/Fiji (NIH, Bethesda, USA) built-in modules, except for images where deconvolution was performed using the softWoRx v4.1.2 built-in module.

To quantify relative promoter strengths, we imaged cells expressing sfGFP from the indicated promoters, using a spinning disk confocal setup comprised of a DMI4000B inverted microscope equipped with an 100x HCX PL APO 6100 (N.A. 1.46) oil objective and Perkin-Elmer Confocal system (including a Yokagawa CSU22 real-time confocal scanning head and solid-state laser lines). Z-series of confocal sections were acquired at 0.4μm intervals using the Volocity software.

For quantifications we prepared an average projection image of three medial Z-sections, Next, we outlined the cell boundaries and measured the mean fluorescence intensity of at least 25 untagged wildtype cells and 25 cells expressing cytosolic sfGFP from indicated promoters. Next, we subtracted the mean fluorescence of wildtype cells from each cell carrying the sfGFP gene that was imaged in identical conditions. We then calculated the mean intensity and standard deviation for cells of the same genotype. Since sfGFP expression levels induced by different promoters varied more than three orders of magnitude, the samples imaged under the same imaging conditions were standardized to the signal from *ura4^+^:p^act1^:sfGFP* cells, which was set to 100 arbitrary units. The data presented is the mean intensity, and standard deviation is denoted.

For images in Figure 4, we concentrated exponentially growing cells by centrifugation at 1000g and spotted them directly between slide and coverslip for immediate imaging. For images in Figure 3 and time-lapse imaging, we placed cells in chambers with solid media made with 2% agarose, as previously described (Vjestica et al., 2016).

### Western blotting and protein level quantification

We grew 70 ml of each strain in EMM media, supplemented with 5 µg/ml of thiamine or 50.25 µg/ml of uracil as indicated, to exponential phase and collected cells by centrifugation for five min at 1000 g. Samples were frozen in liquid nitrogen and stored at −80°C until ready for processing. Subsequent steps were performed at 4°C. Samples were thawed and transferred to 2ml microcentrifuge tubes, washed twice with ice-cold PBS buffer (NaCl 137mM, KCl 2.7mM, Na2HPO4 10mM, KH2PO4 1.8mM, pH7.5, protease inhibitors mix) and re-suspended in 500 µl of the PBS buffer. ~1 ml of acid-washed glass beads were added. Cells were lysed using the FastPrep-24 bead-beater (MP Biomedicals, Santa Ana, USA) set to 4.5 m/s with 10 cycles of 20-second beating and 40 seconds cooling on ice. We then pierced the bottom of the tube with a heated needle, placed that tube into a 1.5ml microfuge tube and centrifuged it at 150 g for 60 s. In this manner, we discarded the beads and transferred the samples to a new tube. Cell debris were pelleted by centrifugation for 15 minutes at 13’000g. We collected the supernatant and determined the protein concentration using the Bradford assay. All samples were adjusted to the same protein concentration and re-suspended in Laemmli buffer, heated to 90°C for 5 min and then subjected to SDS-PAGE (XP10205BOX, Thermo Fischer, Waltham, USA). The gel was then blotted onto nitrocellulose membrane using the Towbin transfer buffer (25mM Tris-base, 192mM glycin, 20% methanol, pH8.3). After blocking in 5% milk, the membrane was first probed with an anti-GFP antibody (Cat.No. 11814460001, Roche, Basel, Switzerland; 1:3’000 dilution) followed by a secondary anti-mouse-IR800 (R-05061, Advansta, Menlo Park, USA), and visualized on the Fusion FX (Vilber, Collégien, France). Subsequently, the same membrane was probed with the TAT-1 antibody (a kind gift from Keith Gull, University of Oxford, UK) targeting tubulin as a loading control. We used ImageJ/Fiji to quantify the signal detected for GFP and normalized it to the tubulin signal. We used the unlabeled area of the membrane as the background intensities. The experiment was qualitatively reproduced in two independent replicates and quantified from the second replicate.

### Flow Cytometry

Flow cytometry was performed on a BD Biosciences Fortessa analyser using CellQuest software. To stain for dead cells in the population, cells were diluted in EMM-ALU medium to final OD 0.4 and 100 μl of cell suspension was mixed with 900 µl EMM-ALU medium containing 1μg/ml propidium iodide. After 30 s of incubation cells were analysed by flow cytometry without gating during acquisition with 10^4^ cells recorded for each sample. Data were analysed using the FlowJo software package using the following gating strategy: a) the main cellular population was distinguished using forward and side scatter to exclude cell aggregates, b) doublets were excluded from analysis by plotting FSC-A versus FSC-H and gating along the diagonal, c) dead cells were excluded from analysis by gating out PI positive events, d) GFP positive events were detected in the green channel. More than 70% of all events passed the indicated criteria.

## Supporting information

Supplemental sequences

## Author contributions

The project was conceived and designed by AV. Experiments were performed by AV and MM. Plasmids were constructed by AV and MM, except pAV0756 and AV0757 that were made by LM, pAV0710 made by GL, pAV0620 made by MB, and pAV0517 made by IBC. PN provided technical assistance. AV, MM and SGM wrote the manuscript.

### Acknowledgements

We are grateful to the Lucie Kesnerova and Philip Engel for help with the qPCR experiments and Serge Pelet for gift of the 3mTagBFP2 construct.

## Data availability

Plasmid and fission yeast strains are available from Addgene and NBRP repositories. All plasmid sequences are made available with the manuscript.

## Supplemental materials

**Table S4**. The qPCR results and analyses shown in the study.

**Movie S1. Timelapse images of cells expressing indicated fluorophores.** Time in hour:min format is indicated at the top of each strain.

**Supplemental Sequences**. Zip compressed file contains sequences annotated in the GenBank format of plasmids and auxotrophic genomic loci generated in the study.

**Source Data D1.** Zip compressed file contains the raw qPCR results in the. eds format used in to generate data in Table S4 and Fig. 1E and 2F.

---

**Figure S1.**
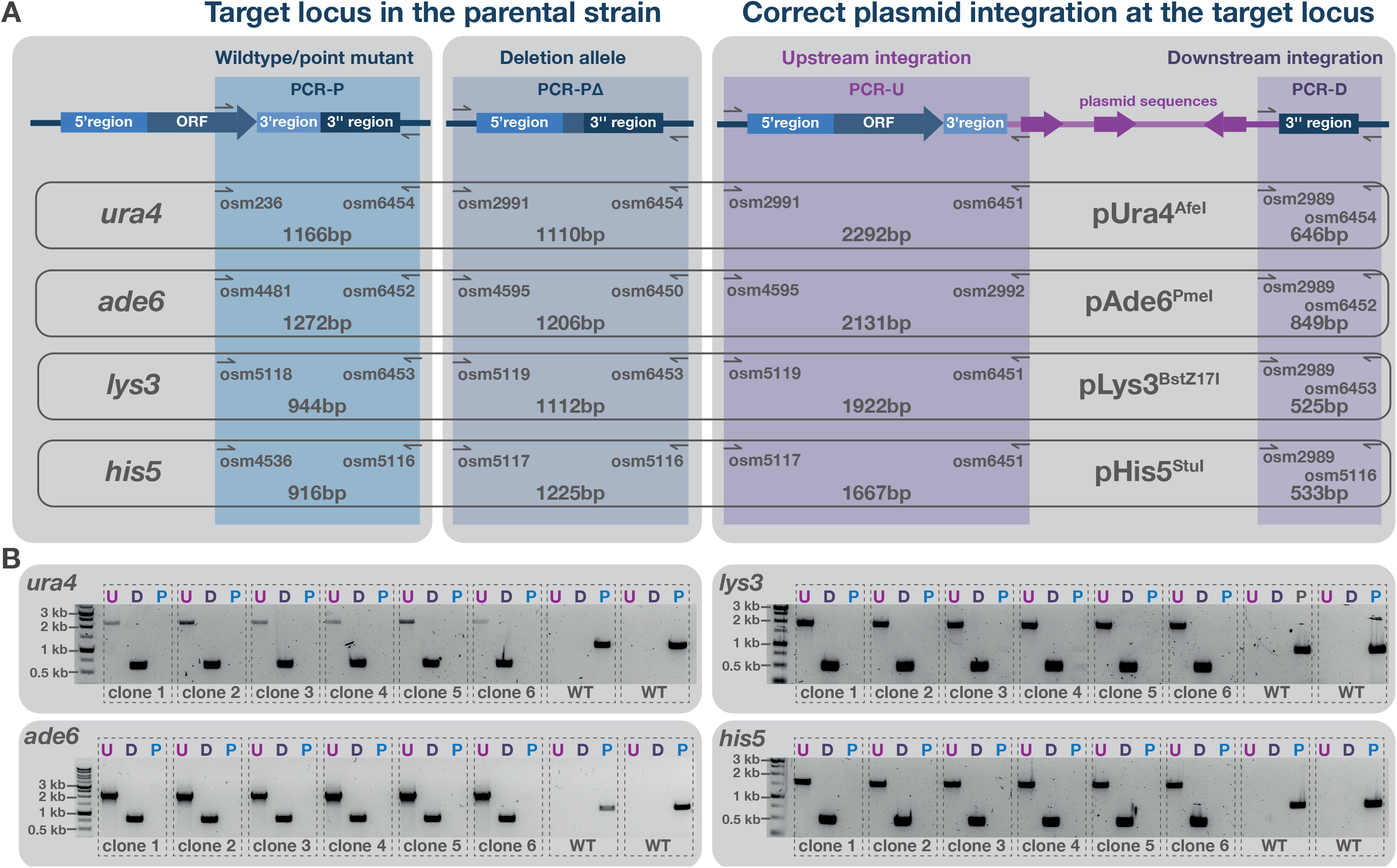
Diagnostic PCRs to test SIV genomic integration. **(A)** Overview of the genotyping strategy to test for correct integration of the plasmids targeting the indicated genomic loci. The top schematic shows parental and transformant loci where arrows denote the primers used for genotyping and colored areas the segment amplified by PCRs. The specific primers carrying the “osm” identifier, and the sizes of the PCR fragments they produce are denoted for each genomic locus. For primer sequences please see Supplemental Table S3. The left and middle panel show the diagnostic PCR for the presence of the parental locus. The PCR-P is used when transformants are obtained from wildtype and point-mutant target alleles, and PCR-P∆ when the parental strain is a deletion of the target locus (*ura4-D18*, *ade6-D19*, *lys3-D20*, *his5-D21*). The right panel shows the diagnostic PCRs to test for the correct upstream (PCR-U) and downstream (PCR-D) integration of the plasmids into target loci. **(B)** The PCR-U, PCR-D and PCR-P results performed on transformants at the indicated loci (clones 1-6) and the parental strain (WT) as detailed in (A).

**Table S1.** List of plasmids used in the study. Please note embedded links to NBRP and Addgene repositories.

| Addgene ID | NBRP ID | Plasmid ID | Plasmid name |
| --- | --- | --- | --- |
| 133467 | FYP4650 | **pAV0133** | pUra4<sup>AfeI</sup> |
| 133468 | FYP4651 | **pAV0356** | pAde6<sup>PmeI</sup> |
| 133469 | FYP4652 | **pAV0357** | pLys3<sup>BstZ17I</sup> |
| 133470 | FYP4653 | **pAV0413** | pHis5<sup>StuI</sup> |
| 133471 | FYP4654 | **pAV0584** | pUra4<sup>AfeI</sup>-natMX |
| 133472 | FYP4655 | **pAV0585** | pAde6<sup>PmeI</sup>-hphMX |
| 133473 | FYP4656 | **pAV0586** | pLys3<sup>BstZ17I</sup>-bsdMX |
| 133474 | FYP4657 | **pAV0587** | pHis5<sup>StuI</sup>-bleMX |
| 133475 | FYP4658 | **pAV0784** | pHis5<sup>StuI</sup>-bsdMX |
| 133476 | FYP4659 | **pAV0661** | pAde6<sup>PmeI</sup>-pact1-sfGFP-terminator<sup>ScCYC1</sup> |
| 133477 | FYP4660 | **pAV0662** | pLys3<sup>BstZ17I</sup>-pact1-sfGFP-terminator<sup>ScCYC1</sup> |
| 133478 | FYP4661 | **pAV0663** | pHis5<sup>StuI</sup>-pact1-sfGFP-terminator<sup>ScCYC1</sup> |
| 133479 | FYP4662 | **pAV0714** | pUra4<sup>AfeI</sup>-pact1-sfGFP-terminator<sup>ScCYC1</sup> |
| 133480 | FYP4663 | **pAV0746** | pUra4<sup>AfeI</sup>-p<sup>rga3</sup>-sfGFP-terminator<sup>ScCYC1</sup> |
| 133481 | FYP4664 | **pAV0747** | pUra4<sup>AfeI</sup>-p<sup>pom1</sup>-sfGFP-terminator<sup>ScCYC1</sup> |
| 133482 | FYP4665 | **pAV0748** | pUra4<sup>AfeI</sup>-p<sup>pak1</sup>-sfGFP-terminator<sup>ScCYC1</sup> |
| 133483 | FYP4666 | **pAV0749** | pUra4<sup>AfeI</sup>-p<sup>tdh1</sup>-sfGFP-terminator<sup>ScCYC1</sup> |
| 133484 | FYP4667 | **pAV0750** | pUra4<sup>AfeI</sup>-p<sup>urg1</sup>-sfGFP-terminator<sup>ScCYC1</sup> |
| 133485 | FYP4668 | **pAV0751** | pUra4<sup>AfeI</sup>-p<sup>nmt1</sup>-sfGFP-terminator<sup>ScCYC1</sup> |
| 133486 | FYP4669 | **pAV0752** | pUra4<sup>AfeI</sup>-p<sup>nmt41</sup>-sfGFP-terminator<sup>ScCYC1</sup> |
| 133487 | FYP4670 | **pAV0753** | pUra4<sup>AfeI</sup>-p<sup>nmt81</sup>-sfGFP-terminator<sup>ScCYC1</sup> |
| 133488 | FYP4671 | **pAV0327** | pUra4<sup>AfeI</sup>-pact1-sfGFP-terminator<sup>tdh1</sup> |
| 133489 | FYP4672 | **pAV0328** | pUra4<sup>AfeI</sup>-pact1-mCherry-terminator<sup>tdh1</sup> |
| 133490 | FYP4673 | **pAV0471** | pUra4<sup>AfeI</sup>-p<sup>tdh1*</sup>-mTagBFP2-terminator<sup>tdh1</sup> |
| 133491 | FYP4674 | **pAV0624** | pUra4<sup>AfeI</sup>-p<sup>tdh1</sup>-3mTagBFP2-terminator<sup>tdh1</sup> |
| 133492 | FYP4675 | **pAV0517** | pHis5<sup>StuI</sup>-p<sup>map3</sup>-GFP-terminator<sup>nmt</sup> |
| 133493 | FYP4676 | **pAV0543** | pLys3<sup>BstZ17I</sup>-p<sup>map3</sup>-mCherry-terminator<sup>ScADH1</sup>-natMX |
| 133494 | FYP4677 | **pAV0761** | pLys3<sup>BstZ17I</sup>-p<sup>map3</sup>-mTagBFP2-terminator<sup>ScADH1</sup>-bleMX |
| 133495 | FYP4678 | **pAV0523** | pAde6<sup>PmeI</sup>-p<sup>mam1*</sup>-sfGFP-terminator<sup>ScADH1</sup>-kanMX |
| 133496 | FYP4679 | **pAV0762** | pAde6<sup>PmeI</sup>-p<sup>mam1*</sup>-mCherry-terminator<sup>ScADH1</sup>-kanMX |
| 133497 | FYP4680 | **pAV0763** | pAde6<sup>PmeI</sup>-p<sup>mam1*</sup>-mTagBFP2-terminator<sup>ScADH1</sup>-kanMX |
| 133498 | FYP4681 | **pAV0478** | pHis5<sup>StuI</sup>-p<sup>tdh1</sup>-NLS<sup>SV40</sup>-sfGFP-NLS<sup>SV40</sup>-natMX |
| 133499 | FYP4682 | **pAV0479** | pHis5<sup>StuI</sup>-p<sup>tdh1</sup>-NES<sup>wis1</sup>-sfGFP-NES<sup>mia1</sup>-natMX |
| 133500 | FYP4683 | **pAV0526** | pHis5<sup>StuI</sup>-p<sup>tdh1</sup>-sfGFP-NLS<sup>SV40</sup>-natMX |
| 133501 | FYP4684 | **pAV0765** | pLys3<sup>BstZ17I</sup>-p<sup>tdh1*</sup>-NLS<sup>SV40</sup>-mCherry-terminator<sup>ScADH1</sup>-kanMX |
| 133502 | FYP4685 | **pAV0532** | pLys3<sup>BstZ17I</sup>-p<sup>tdh1*</sup>-NLS<sup>SV40</sup>-mTagBFP2-terminator<sup>tdh1</sup>-kanMX |

Table S1 continued.
| Addgene ID | NBRP ID | Plasmid ID | Plasmid name |
| --- | --- | --- | --- |
| 133503 | FYP4686 | **pAV0605** | pUra4<sup>AfeI</sup>-p<sup>act1</sup>-sfGFP-RitC-terminator<sup>tdh1</sup> |
| 133504 | FYP4687 | **pAV0607** | pAde6<sup>PmeI</sup>-p<sup>act1</sup>-mCherry-RitC-terminator<sup>ScADH1</sup> |
| 133505 | FYP4688 | **pAV0612** | pAde6<sup>PmeI</sup>-p<sup>tdh1</sup>-3mTagBFP2-RitC-terminator<sup>ScADH1</sup>-bsdMX |
| 133506 | FYP4689 | **pAV0620** | pUra4<sup>AfeI</sup>-p<sup>pcn1</sup>-eGFP-pcn1-terminator<sup>nmt1</sup>-natMX |
| 133507 | FYP4690 | **pAV0785** | pUra4<sup>AfeI</sup>-p<sup>pcn1</sup>-mCherry-pcn1-terminator<sup>nmt1</sup>-natMX |
| 133508 | FYP4691 | **pAV0756** | pHis5<sup>StuI</sup>-p<sup>act1</sup>-CRIB(gic2<sup>aa2-181</sup>)-3mCherry-bsdMX |
| 133509 | FYP4692 | **pAV0757** | pHis5<sup>StuI</sup>-p<sup>pak1</sup>-CRIB(gic2<sup>aa2-181</sup>)-3GFP-terminator<sup>ScADH1</sup>-kanMX |
| 133510 | FYP4693 | **pAV0816** | pAde6<sup>PmeI</sup>-p<sup>act1</sup>-CRIB(gic2<sup>aa2-181</sup>)-mTagBFP2-terminator<sup>ScADH1</sup>-bsdMX |
| 133511 | FYP4694 | **pAV0758** | pLys3<sup>BstZ17I</sup>-p<sup>pak1</sup>-RasAct(3xbyr2RBD)-3GFP-kanMX-terminator<sup>ScADH1</sup> |
| 133512 | FYP4695 | **pAV0835** | pUra4<sup>AfeI</sup>-p<sup>act1</sup>-mCherry-CHD<sup>Rng2</sup>-terminator<sup>ScCYC1</sup> |
| 133513 | FYP4696 | **pAV0771** | pAde6<sup>PmeI</sup>-p<sup>act1</sup>-3mTagBFP2-CHD<sup>Rng2</sup>-terminator<sup>ScCYC1</sup> |
| 133514 | FYP4697 | **pAV0782** | pAde6<sup>PmeI</sup>-p<sup>act1</sup>-LifeAct-sfGFP-terminator<sup>ScADH1</sup>-bsdMX |
| 133515 | FYP4698 | **pAV0783** | pAde6<sup>PmeI</sup>-p<sup>act1</sup>-LifeAct-mCherry-terminator<sup>ScADH1</sup>-bsdMX |
| 133516 | FYP4699 | **pAV0772** | pAde6<sup>PmeI</sup>-p<sup>atb2</sup>-sfGFP-atb2-terminator<sup>atb2</sup>-hphMX |
| 133517 | FYP4700 | **pAV0710** | pAde6<sup>PmeI</sup>-p<sup>atb2</sup>-mCherry-atb2-terminator<sup>atb2</sup>-hphMX |
| 133518 | FYP4701 | **pAV0770** | pAde6<sup>PmeI</sup>-p<sup>atb2</sup>-mTagBFP2-atb2-terminator<sup>atb2</sup>-hphMX |
| 133519 | FYP4702 | **pAV0773** | pLys3<sup>BstZ17I</sup>-p<sup>BiP</sup>-SignalSequence<sup>BiP</sup>-sfGFP-AHDL-bsdMX |
| 133520 | FYP4703 | **pAV0764** | pLys3<sup>BstZ17I</sup>-p<sup>BiP</sup>-SignalSequence<sup>BiP</sup>-mCherry-AHDL-bsdMX |
| Not deposited | Not deposited | **pAV0569** | pUra4<sup>AfeI</sup>-p<sup>tdh1</sup>*-sfGFP-terminator<sup>tdh1</sup> |
| Not deposited | Not deposited | **pAV0570** | pAde6<sup>PmeI</sup>-p<sup>tdh1</sup>*-sfGFP-terminator<sup>tdh1</sup> |
| Not deposited | Not deposited | **pAV0571** | pLys3<sup>BstZ17I</sup>-p<sup>tdh1</sup>*-sfGFP-terminator<sup>tdh1</sup> |
| Not deposited | Not deposited | **pAV0572** | pHis5<sup>StuI</sup>-p<sup>tdh1</sup>*-sfGFP-terminator<sup>tdh1</sup> |
| Not deposited | Not deposited | **pAV0596** | pAde6<sup>PmeI</sup>-ura4Casette |
| Not deposited | Not deposited | **pAV0597** | pLys3<sup>BstZ17I</sup>-ura4Casette |
| Not deposited | Not deposited | **pAV0598** | pHis5<sup>StuI</sup>-ura4Casette |
| Not deposited | Not deposited | **pAV0616** | pAde6<sup>PmeI</sup>-ura4casette-hphMX |
| Not deposited | Not deposited | **pAV0617** | pLys3<sup>BstZ17I</sup>-ura4casette-bsdMX |
| Not deposited | Not deposited | **pAV0618** | pHis5<sup>StuI</sup>-ura4casette-bleMX |
| Not deposited | Not deposited | **pAV0623** | pJK210-natMX |

**Table S2.**
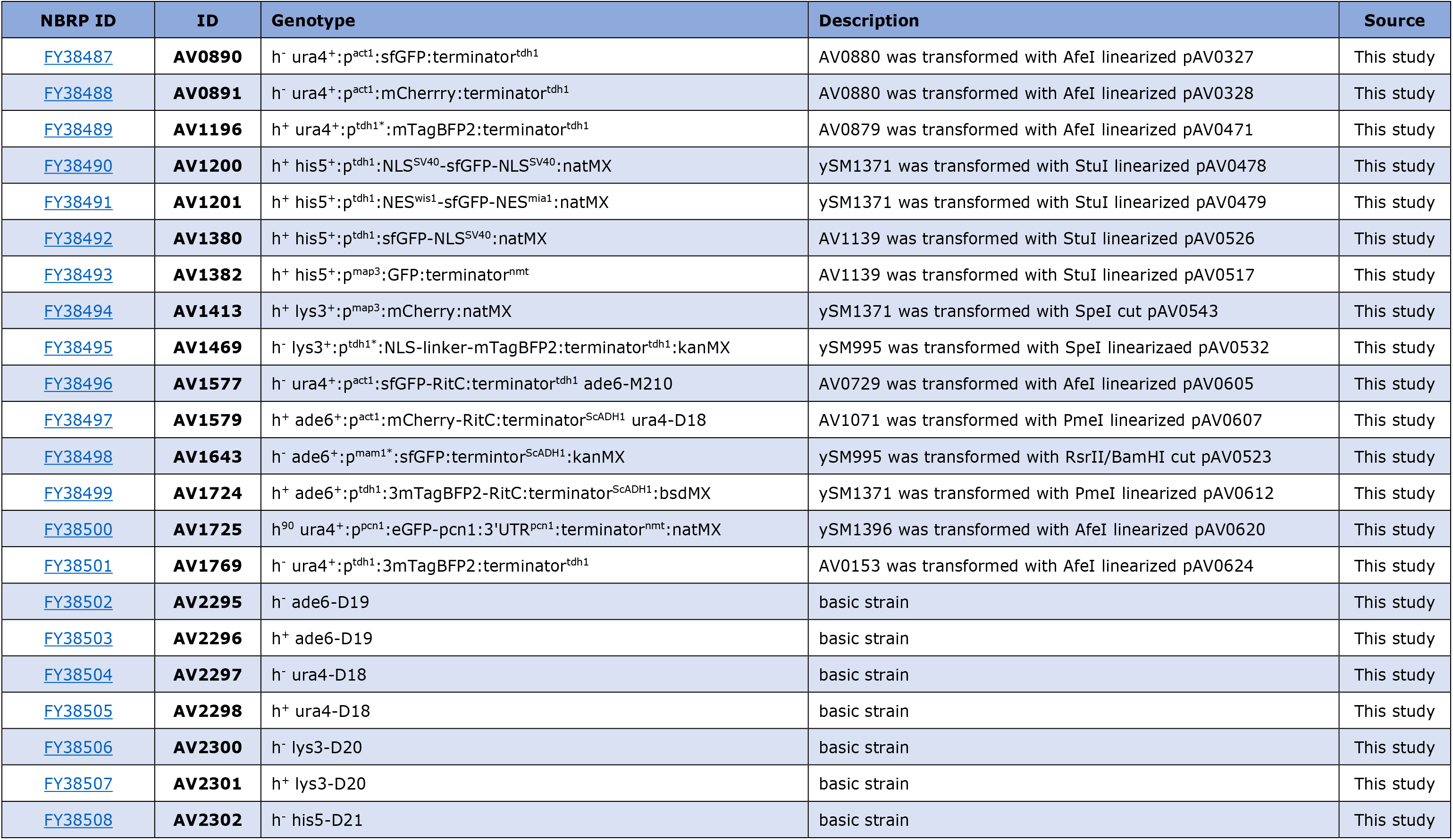
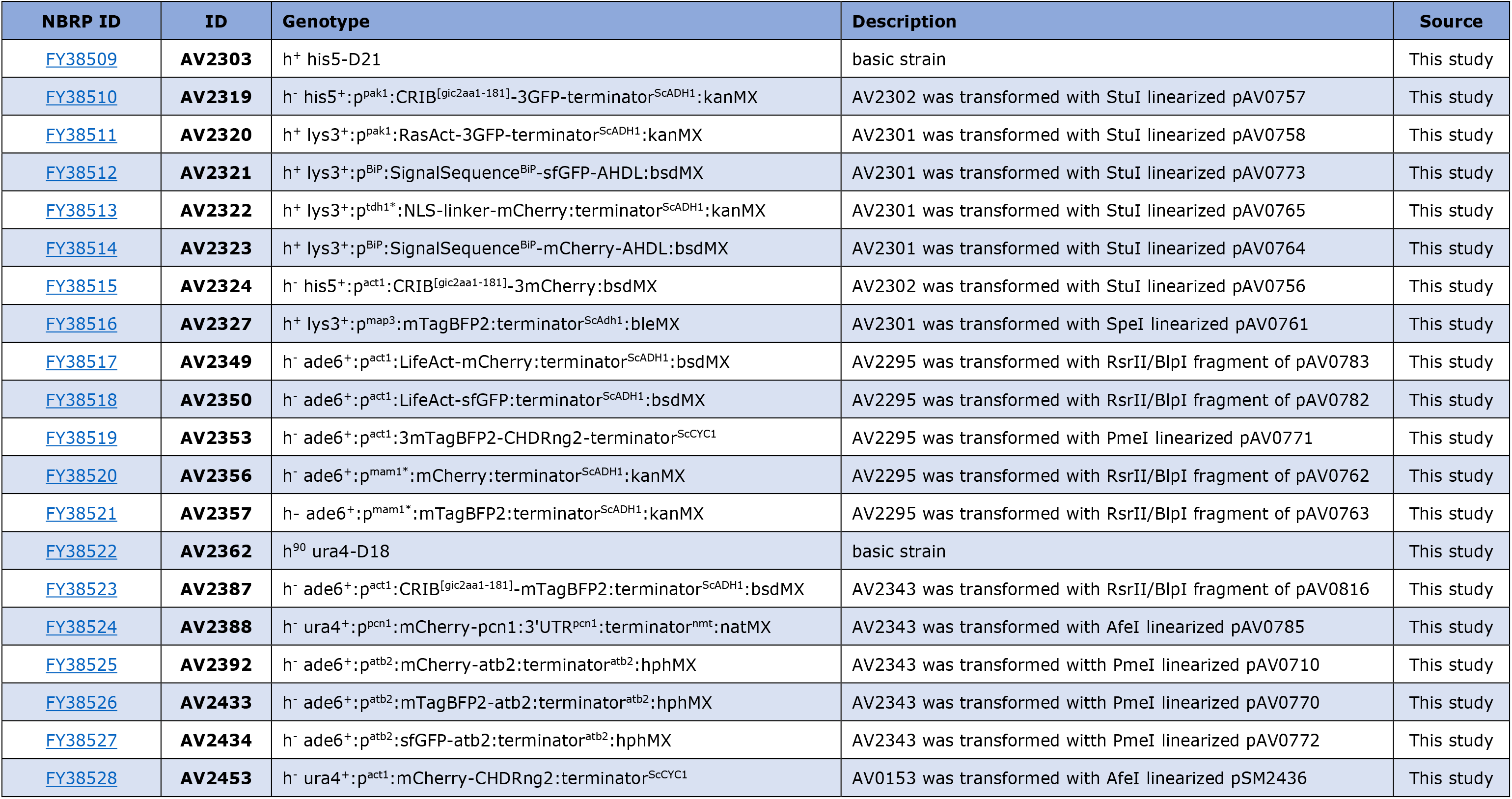
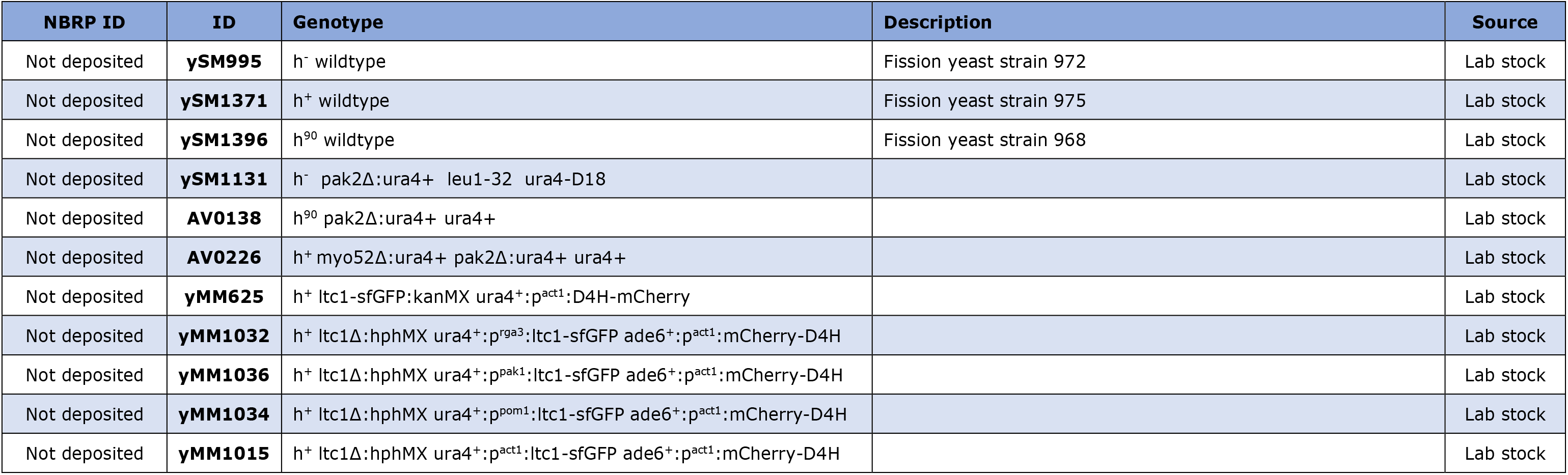
List of fission yeast strains used in the study. Please note embedded links to NBRP repository.

**Table S3.**
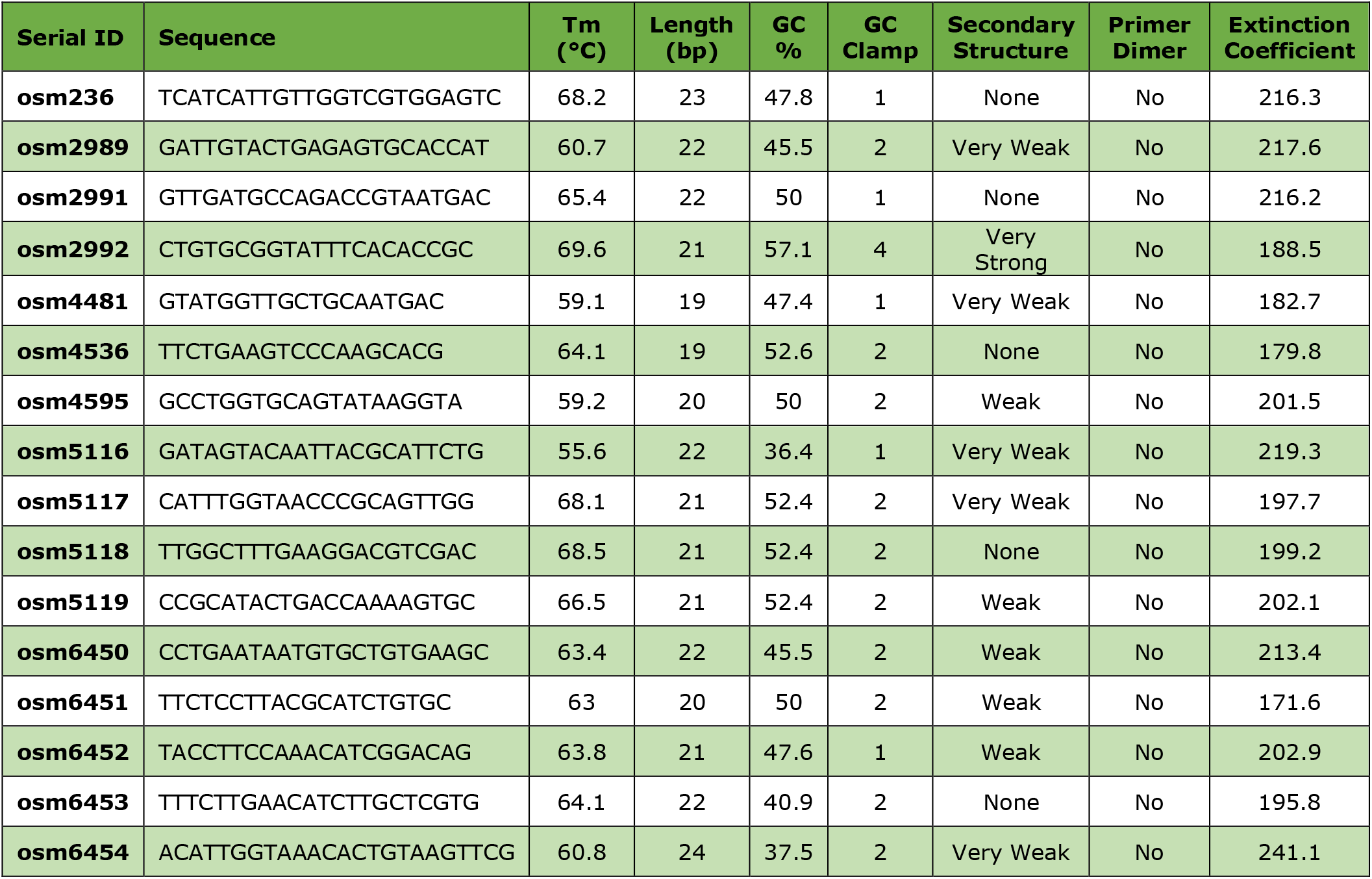
List of primers used for genotyping. See Figure S1 for usage.

